# An *in vitro* vesicle formation assay to analyze protein sorting in the secretory transport pathway

**DOI:** 10.1101/2020.02.12.945162

**Authors:** Yan Huang, Haidi Yin, Xiao Tang, Qian Wu, Mo Wang, Kristina Poljak, Baiying Li, Liu Yang, Zhixiao Wu, Liwen Jiang, Elizabeth A. Miller, Zhong-Ping Yao, Yusong Guo

## Abstract

The fidelity of protein transport in the secretory transport pathway relies on the accurate sorting of proteins to their correct destination. To deepen our understanding of the underlying molecular mechanisms, it is important to develop a robust approach to systematically reveal cargo proteins that depend on a specific cargo sorting machinery to be efficiently packaged into vesicles. Here, we used an *in vitro* assay that reconstitutes packaging of human cargo proteins into vesicles to quantify cargo capture. Quantitative mass spectrometry analyses of the isolated vesicles revealed novel cytosolic proteins that are associated with vesicle membranes in a GTP-dependent manner or that interact with GTP-bound Sar1A on vesicle membranes. Functional analysis indicates that two of them, FAM84B and PRRC1, regulate anterograde trafficking. Comparing control cells with cells depleted of the cargo receptors, SURF4 or ERGIC53, we revealed specific clients of each of these two export adaptors. Moreover, our results indicate that vesicles enriched with a specific cargo protein contain specific transmembrane cargo and SNARE proteins. A SNARE protein, Vti1B, is identified to be in vesicles enriched with a planar cell polarity protein, Frizzled6, and promotes vesicular release of Frizzled6. Our results indicate that the vesicle formation assay in combination with quantitative mass spectrometric analysis is a robust and powerful tool to reveal novel cytosolic and transmembrane proteins that regulate trafficking of a specific cargo protein.

## Introduction

The eukaryotic secretory pathway plays important roles in delivering a variety of newly synthesized proteins to their specific resident compartments. The fidelity of protein transport in the secretory pathway depends on accurate sorting of specific cargo proteins into transport vesicles. Defects in cargo sorting cause protein mistargeting and induce defects in establishing cell polarity, immunity as well as other physiological processes (Guo et al., 2014).

A variety of cytosolic proteins are recruited to the membrane and play important roles in the protein sorting process. These cytosolic proteins include small GTPases of the Arf family and cargo adaptors (Guo et al., 2014; Lee et al., 2004). The Arf family GTPases cycle between a GDP-bound cytosolic state and a GTP-bound state. Upon GTP binding, Arf proteins undergo conformational changes in which the N-terminal amphipathic helix is exposed to bind membranes and the switch domains change their conformation to recruit various cytosolic cargo adaptors. Once recruited onto the membranes, these cargo adaptors recognize sorting motifs on the cargo proteins. This recognition step is important for efficiently capturing cargo proteins into vesicles.

The Arf family protein, Sar1, regulates packaging of cargo proteins into vesicles at the endoplasmic reticulum (ER). GTP-bound Sar1 mediates membrane recruitment of the coat protein complex II (COPII) to capture cargo proteins (Lee et al., 2004). Soluble cargo proteins in the lumen of the ER cannot be directly recognized by COPII coat and those proteins are thought to be linked to the cargo sorting machinery on the cytosolic side by transmembrane cargo receptors. One cargo receptor in mammalian cells, ERGIC53, is a mannose lectin and functions in capturing specific N-linked glycoproteins in the lumen of the ER (Dancourt and Barlowe, 2010). ERGIC53 regulates ER export of blood coagulation factors V and VIII, a cathepsin-Z-related protein and alpha1-antittrypsin (Appenzeller et al., 1999; Nichols et al., 1998; Nyfeler et al., 2008; Vollenweider et al., 1998). The p24 family proteins function as cargo receptors to regulate ER export of specific GPI-anchored proteins in mammalian cells (Barlowe and Helenius, 2016). The mammalian homologs of yeast ER vesicle (Erv) proteins have also been thought to function as cargo receptors (Dancourt and Barlowe, 2010). One of these proteins, SURF4, binds amino-terminal tripeptide motifs of soluble cargo proteins and regulates ER export of soluble cargo proteins including the yolk protein VIT-2 in Caenorhabditis elegans (Saegusa et al., 2018), and PCSK9 and apolipoprotein B in mammalian cells (Emmer et al., 2018; Yin et al., 2018).

To deepen our understanding of the protein sorting process in the secretory pathway, it is important to utilize a robust approach to systematically reveal cargo proteins that depend on a specific factor to be efficiently packaged into vesicles. Revealing this will provide significant insight into the functions and the specificity of the cargo sorting process. Since distinct cytosolic proteins are recruited to membranes by different GTP-bound Arf family proteins, systematic approaches are needed to characterize budding events associated with a specific GTP-bound Arf family protein. Moreover, a transport vesicle should not only contain the correct cargo molecules but also the correct SNAREs and motor proteins to ensure targeting of vesicles to the correct destination. Some cargo proteins rely on cargo receptors or other cargo proteins to be recognized by the sorting machinery. Thus a better understanding of the protein composition of vesicles containing specific cargo proteins will provide novel information about the vesicular trafficking process.

A microscope-based approach, PAIRS, has been utilized to identify the spectrum of cargo proteins that depends on a specific cargo receptor for the ER export process in yeast. This analysis is focused on around 150 cargo molecules with fluorescent tags (Herzig et al., 2012). An *in vitro* assay that reconstitutes packaging of cargo proteins into vesicles has been used to reveal protein profiles of vesicles budded with purified COPII or COPI proteins (Adolf et al., 2019). However, this analysis did not identify any non-ER resident transmembrane proteins or secretory proteins (Adolf et al., 2019). This is possibly due to an unappreciated requirement for other cytosolic factors in addition to the COP coats. Affinity chromatography has been utilized to reveal cytosolic proteins that specifically interact with GTP bound Arf or Rab proteins (Christoforidis and Zerial, 2000; Guo et al., 2013; Jin et al., 2010). In this approach, the membranes are disrupted, which might preclude identification of membrane-associated effectors. Thus, it is important to develop additional approaches to reveal novel cytosolic proteins that associate with GTP-bound Arf proteins on membranes.

Here, we used an in *vitro* assay to reconstitute packaging of cargo proteins into transport vesicles utilizing rat liver cytosol as source of cytosolic proteins. Analysis of vesicle fractions by quantitative mass spectrometry analysis revealed novel cytosolic proteins that are associated with vesicles dependent on GTP or GTP-bound Sar1A and that regulate protein trafficking in the secretory transport pathway. We also revealed cargo proteins that depend on a specific cargo receptor, ERGIC53 or SURF4, to be efficiently packaged into vesicles. Moreover, we revealed the specificity of cargo packaging, identifying proteins, including SNAREs, that are selectively associated with a subset of cargo proteins, EGFR or Frizzled6, in transport vesicles. Our study indicates that the vesicle formation assay is a robust tool to reveal functional roles of specific factors in protein sorting, and to reveal novel factors that regulate vesicular trafficking.

## Results

### An *in vitro* reconstituted vesicle formation assay for proteomic analysis

An *in vitro* vesicle formation assay to reconstitute packaging of cargo proteins into vesicles from mammalian cells has been well-established (Kim et al., 2007; Ma et al., 2018; Merte et al., 2010; Niu et al., 2019). We sought to perform this assay in HEK293T cells on a large scale and then perform proteomic analysis on the isolated vesicles. The general procedures of the vesicle formation assay are shown in Figure 1A. Briefly, HEK293T cells were permeabilized by digitonin, after which the semi-intact cells were washed with buffer to remove cytosolic proteins. Washed semi-intact cells were then incubated at 30°C with rat liver cytosol, GTP and an ATP regeneration system (ATPrS). The small vesicles released during this incubation were separated from the heavy donor membranes by medium speed centrifugation. The supernatant containing the vesicle fraction was adjusted to 35% Opti-Prep and overlaid with layers of 30% Opti-Prep and the reaction buffer. The samples were then centrifuged to float the vesicles away from cytosolic proteins that are not associated with membranes. Two control experiments were performed: one performed in the absence of GTP and ATPrS and the other performed in the presence of a non-hydrolysable analog of GTP, GMPPNP.

**Figure 1.**
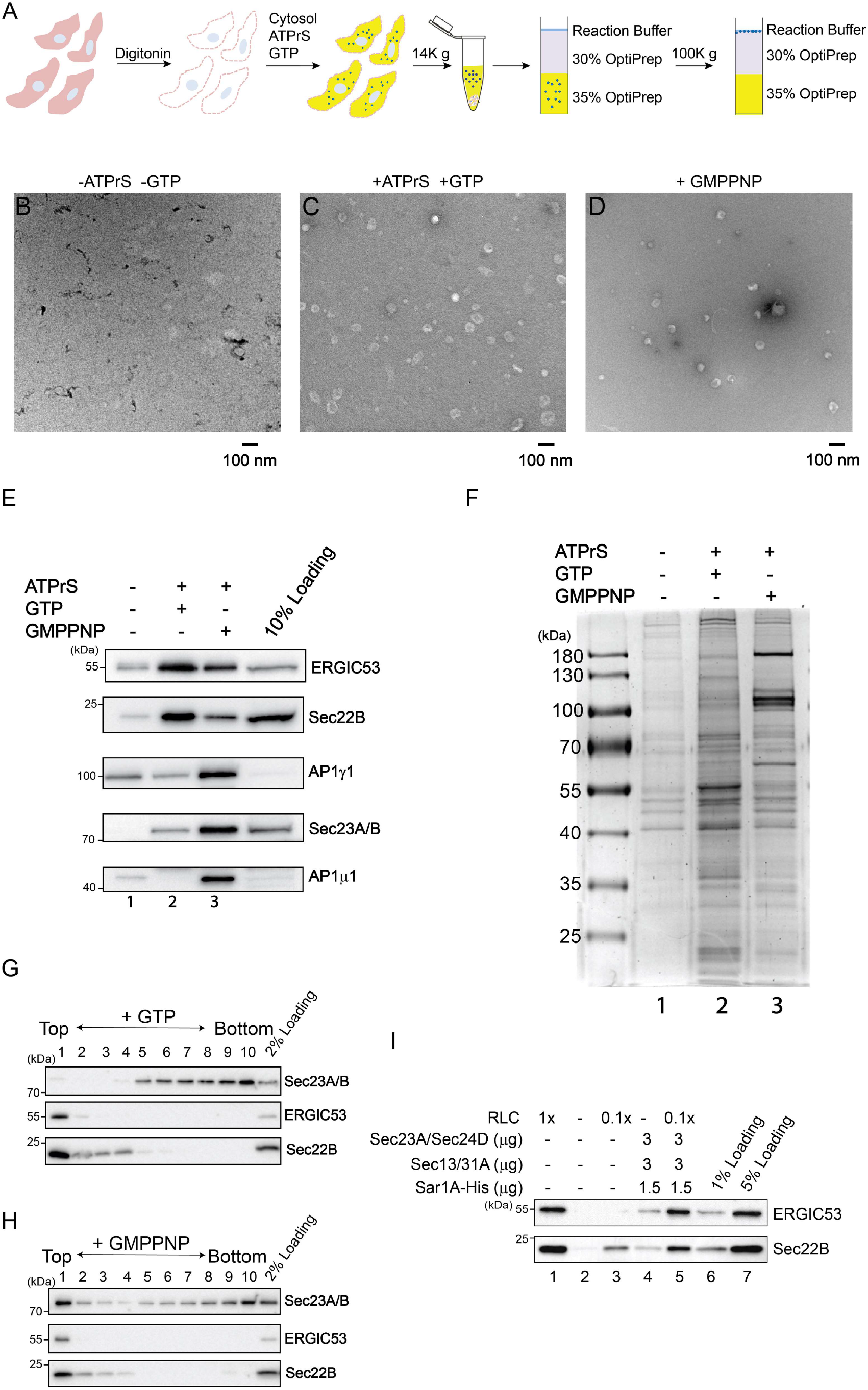
A large scale *in vitro* formation assay for proteomic analysis. **A.** Diagram demonstrating the experimental procedures for the vesicle formation assay. **B-D.** Visualization of the morphology of the buoyant membrane structures formed in the budding reaction. The buoyant membranes were isolated by density gradient flotation and analyzed by negative stain EM. *Scale bar*, 100 nm. **E-F.** The vesicle formation assay was performed using the indicated reagents. Vesicle fractions were analyzed by immunoblot (E) or Coomassie blue staining (F). ATPrS: ATP regeneration system. **G-H.** The vesicle formation assay was performed in the presence of GTP (G) or GMPPNP (H). The vesicle fractions were evaluated by density gradient flotation. **I.** Vesicle formation assay was performed using the indicated reagents. The vesicle fraction was analyzed by immunoblot using the indicated antibodies.

We performed negative stain electron microscopy to visualize the morphology of the buoyant membrane structures produced in the vesicle budding reaction. We detected numerous small membrane structures (Figures 1C) with an average diameter of 53 ± 18 nm. When we performed the vesicle budding reaction in the absence of GTP and an ATPrS or in the presence of GMPPNP, the number of vesicles was greatly reduced (Figure 1B, 1D). These analyses are consistent with a previous report (Ma et al., 2018) and are consistent with the slowly-sedimenting membranes in the budding reaction representing transport vesicles rather than fragments of the ER or Golgi.

The buoyant membranes were analyzed by immunoblotting with antibodies against standard cargo proteins in COPII vesicles, Sec22B (a tSNARE) and ERGIC53. Capture of ERGIC53 and Sec22B into the vesicle fraction was enhanced by the ATPrS and GTP (Figure 1E, compare lanes 1 and 2), and was reduced in the presence of GMPPNP (Figure 1E, compare lanes 2 and 3), suggesting GTP hydrolysis is important for efficient packaging of cargo proteins into transport vesicles. In contrast, vesicle coat proteins, including the γ and μ subunit of the adaptor complex 1 (AP-1) and the COPII subunit Sec23A/B, were more robustly associated with the vesicle fraction in the presence of GMPPNP (Figure 1E, compare lanes 2 and 3). These results confirm that GTP hydrolysis permits release of AP-1 and COPII from membranes (Guo et al., 2014; Lee et al., 2004), and that this recycling is important to sustain efficient vesicle formation. We next analyzed the proteins in the buoyant vesicle fractions by SDS-PAGE and Coomassie blue staining (Figure 1F), noting distinct protein complements for the different reaction conditions. Again, the pattern of protein recovery is consistent with coat proteins stabilized in the presence of GMPPNP, and more robust vesicle release in the context of ATPrS and GTP. Finally, we assessed the distribution of cargo and coat proteins throughout the OptiPrep gradient, finding Sec22B and ERGIC53 enriched in the top fraction (Figure 1G-H). We detected Sec23A/B in the floated fraction only when the vesicle formation assay was performed in the presence of GMPPNP (Figure 1G-H)

Since cytosol was used as the source of coat proteins in these experiments, multiple different types of vesicles may be formed. We tested whether purified COPII proteins could promote vesicular release of ER-Golgi cargo proteins. Indeed, Sec22B and ERGIC53 were packaged into the vesicle fraction with purified COPII, albeit with reduced efficiency compared to reactions with cytosol (Figure 1I, compare lanes 1, 2 and 4). Interestingly, when rat liver cytosol at low concentration was supplemented with purified COPII, release of Sec22B and ERGIC53 into vesicles was enhanced (Figure 1I, compare lane 5 with lanes 3 and 4). This result indicates that some proteins in rat liver cytosol work together with purified COPII to promote packaging of cargo proteins into vesicles. Thus, we utilized cytosol prepared from rat liver to provide the source of cytosolic proteins in the vesicle formation assay for our subsequent quantitative analysis.

### Identification of cytosolic proteins specifically associated with vesicles in a GTP-dependent manner

Our results indicate that inhibiting GTP hydrolysis locks the vesicle coat on membranes, presumably by stabilizing interactions with GTP-bound Arf family proteins. To gain a comprehensive view of cytosolic proteins that are associated with vesicle membranes in a GTP-dependent manner, we performed label-free quantitative mass spectrometry to compare protein profiles of the vesicle fractions in GTP vs. GMPPNP treatment conditions based on 3 biological repeats. A total of 1285 proteins were identified and quantified, all of which had two or more unique peptides (FDR<0.01) and were successfully quantified in all of the three biological repeats (Table S1, sheet 1). The fold changes in the identified proteins in the GMPPNP group compared with the GTP group were quantified. Based on protein abundance, a p-value was calculated and plotted against the mean log2 fold changes. Through this quantification approach, 58 proteins were identified as having more than 2-fold enrichment in the GMPPNP group over the GTP group (p < 0.05, Figure 2A, Area B, proteins identified using the protein sequence database of *Homo sapiens* are indicated in round shapes and additional proteins identified using database of *Rattus norvegicus* are indicated in triangle shapes, Table S1, sheet 2). 40 proteins (69%) were known Arf family proteins, Rab proteins and cargo adaptors.

**Figure 2.**
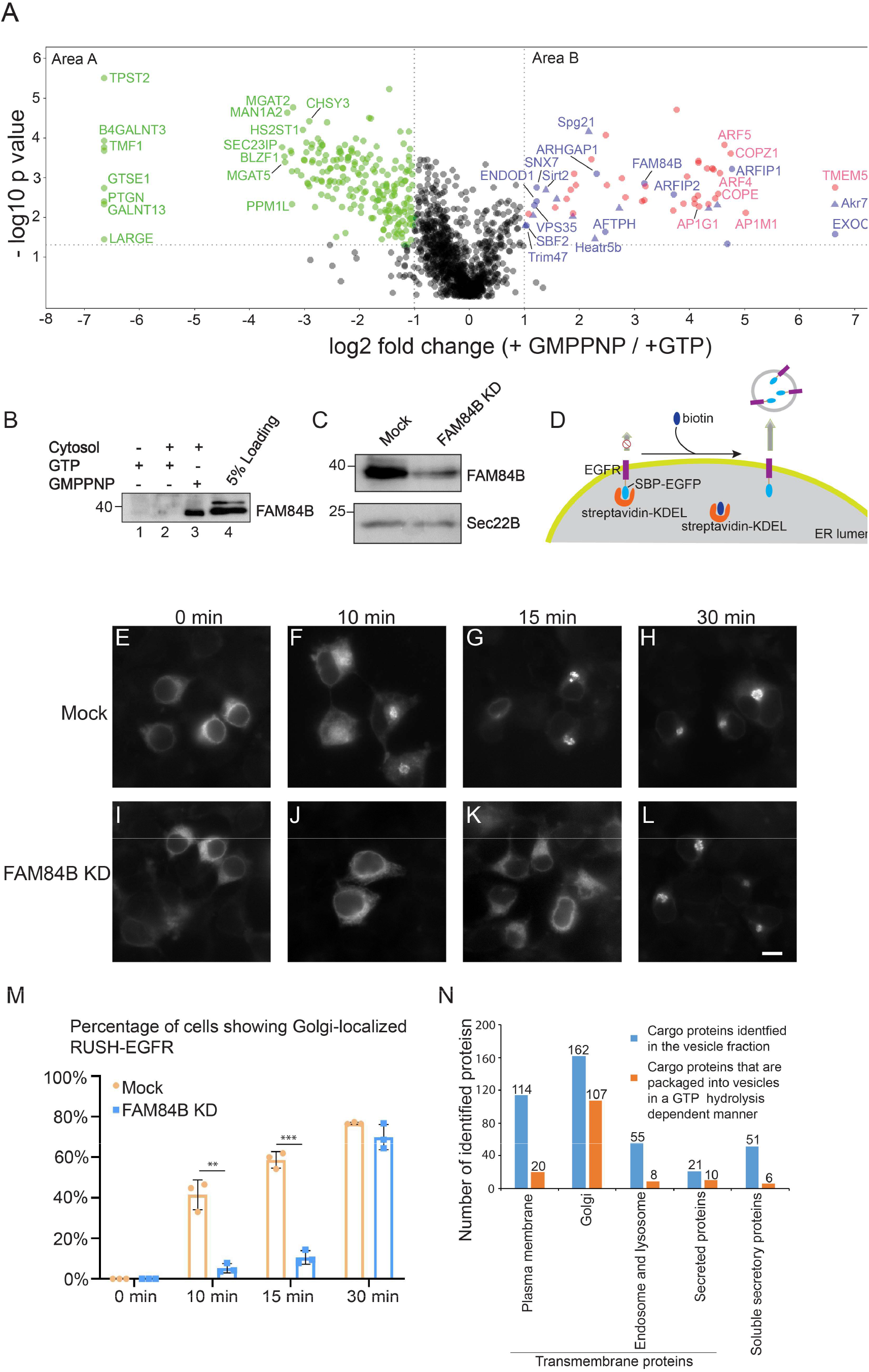
Identification of cytosolic proteins that are associated with vesicles in a GTP dependent manner and cargo proteins that are packaged into vesicles in a GTP-hydrolysis dependent manner. **A.** The vesicle formation assay was performed in the presence of GTP or GMPPNP. The isolated vesicles in each experimental group were resuspended in RapiGest SF surfactant. The proteins in the vesicle fractions were trypsin digested and analyzed by label-free mass spectrometry. A total of 1285 proteins were identified in both experimental groups. The log2 ratio of the abundance of each identified protein in the vesicles prepared in the presence of GMPPNP over that in the vesicles prepared in the presence of GTP was plotted on the x axis and the −log10 p-value of the difference was plotted on the y-axis. **B.** Vesicle formation assay was performed using the indicated regents. The proteins in the vesicle fraction was analyzed by western blot. **C.** HEK293T cells were transfected with control siRNA or siRNA against FAM84B. Day 3 after transfection, cells were lysed and analyzed by western blot. **D.** Diagram demonstrating the RUSH assay. **E-L.** HEK293T cells were transfected with control siRNA (E-H) or siRNA against FAM84B (I-L). 24hr after transfection, cells were re-transfected with plasmids encoding both EGFR^RUSH^ and Str-KDEL. On day 3 after knockdown, cells were incubated with biotin and cycloheximide for the indicated time and the localization of EGFR^RUSH^ was analyzed by fluorescent microscope. Scale bar, 10 μm. **M.** Quantification of the percentage of cells showing Golgi-localized EGFR^RUSH^ in cells treated with control siRNA or siRNA against FAM84B (mean ± S.D.; n = 3; >100 cells counted for each experiment). **, p<0.01; ***, p < 0.001. **N.** Number of proteins categorized based on predictions from Uniprot.

We identified several cytosolic proteins in addition to Arf, Rab and known cargo adaptor proteins that are associated with vesicles in a GTP-dependent manner (Figure 2A Area B, marked in Blue, Table S1, sheet 4). We hypothesize that these proteins may be novel cargo adaptors or proteins associated with vesicle coats. Western blot analysis confirmed that one of these proteins, FAM84B, was significantly enhanced when the incubation was conducted in the presence of GMPPNP (Figure 2B, compare lanes 2 and 3). FLAG-tagged FAM84B colocalized with an ER marker, protein disulfideisomerase (PDI) (Figure S1A-C). In some of the expressing cells, FAM84B-FLAG was partially located at the juxtanuclear Golgi area (Figure S1D-F). We then analyzed the role of FAM84B in the anterograde trafficking along the secretory pathway. We selected epidermal growth factor receptor (EGFR) as a cargo protein and analyzed trafficking of EGFR through a Retention Using Selective Hook (RUSH) transport assay (Boncompain et al., 2012; Mao et al., 2019). In the RUSH assay, HEK293T cells were transfected with a plasmid encoding human EGFR tagged with EGFP and the streptavidin binding peptide (SBP) (EGFR^RUSH^). This plasmid also encodes streptavidin fused to a C-terminal ER retention signal (Lys-Asp-Glu-Leu; Str-KDEL). Due to the binding between streptavidin and SBP, EGFR^RUSH^ was retained at the ER (Figure 2D-E and 2I, 0 min) and colocalized with FAM84B-FLAG (Figure S1G-I). When cells were incubated with biotin, SBP is released from streptavidin thereby releasing EGFR^RUSH^ from the ER (Figure 2D-H, quantification in Figure 2M). Knockdown of FAM84B causes a significant delay of EGFR transport from the ER to the Golgi in the RUSH transport system (Figure 2C, I-L, quantification in Figure 2M) indicating that FAM84B is important for ER-to-Golgi transport of EGFR.

### Classification of the cargo proteins that are packaged into vesicles in the vesicle formation assay

Next, we classified the cargo proteins that are packaged into vesicles in our vesicle formation assay into two groups: 1) soluble secretory cargo proteins; 2) transmembrane proteins that are localized at the Golgi, endosomes, lysosomes or plasma membrane. 4% (51 proteins) of the proteins identified in the vesicle fraction were predicted by Uniprot annotation to be soluble secretory cargo proteins (Figure 2N, Table S1, sheet 5). 37% (480 proteins, Table S1, sheet 5) of the proteins identified in the vesicle fraction were predicted to be transmembrane proteins. Among the predicted transmembrane proteins, 162 of them are predicted to show Golgi localization, 21 proteins can be secreted presumably in extracellular vesicles, 114 proteins are predicted to show plasma membrane localization and 55 proteins show endosomal and lysosomal localization (Figure 2N, Table S1, sheet 5).

Our analyses indicate that the abundances of certain cargo proteins are more enriched in the vesicle fraction when the vesicle formation assay is performed in the presence of GTP than in the presence of GMPPNP. We found that 217 proteins showed more than 2-fold enrichment in the GTP group over the GMPPNP group (p < 0.05, Figure 2A, Area A, Table S1, sheet 3). 72% (156 in total) proteins among the identified proteins in Area A are predicted by Uniprot to be transmembrane proteins: 109 of them are predicted to show Golgi localization, 10 of the predicted transmembrane proteins can be secreted, 20 proteins are predicted to show plasma membrane localization and 8 show endosomal and lysosomal localization (Figure 2N, Table S1, sheet 6). 6 of the identified proteins in Area A are soluble secretory cargo proteins (Figure 2N). We propose that these transmembrane proteins and soluble secretory proteins are cargo proteins that are packaged into vesicles in a GTP hydrolysis dependent manner.

### Identification of cargo proteins and cytosolic proteins that are dependent on Sar1A to be associated with transport vesicles

Next, we sought to utilize this assay to identify cytosolic proteins and cargo proteins that depend on a specific Arf family protein to be incorporated into transport vesicles. We focused our analysis on the ER Arf family member, Sar1, which initiates assembly of the COPII coat (Lee et al., 2004). Sar1 has two isoforms in mammalian cells: Sar1A and Sar1B (Lee et al., 2004). The H79G mutation locks Sar1A in its GTP-bound form and inhibits the COPII-dependent ER export process (Aridor et al., 1995). Consistent with previous reports, Sar1A(H79G) significantly abolished the vesicular capture of Sec22B and ERGIC53 (Figure 3A). In contrast, Sar1A(H79G) did not interfere with the vesicular release of TGN46 (Figure 3B), a cargo protein that cycles between the plasma membrane and the Golgi. Moreover, we found that Sar1A(H79G) enhanced membrane association of the COPII subunit, Sec23A (Figure 3C, compare lanes 3 and 2). In contrast, the dominant active form of another small GTPase, Arfrp1(Q79L), did not enhance the membrane association of Sec23A (Figure 3C, compare lanes 4 and 3). These analyses suggest that our vesicle formation assay recapitulates the specific functions of Sar1A in meditating assembly of COPII coat proteins and in regulating packaging of cargo proteins in COPII vesicles.

**Figure 3.**
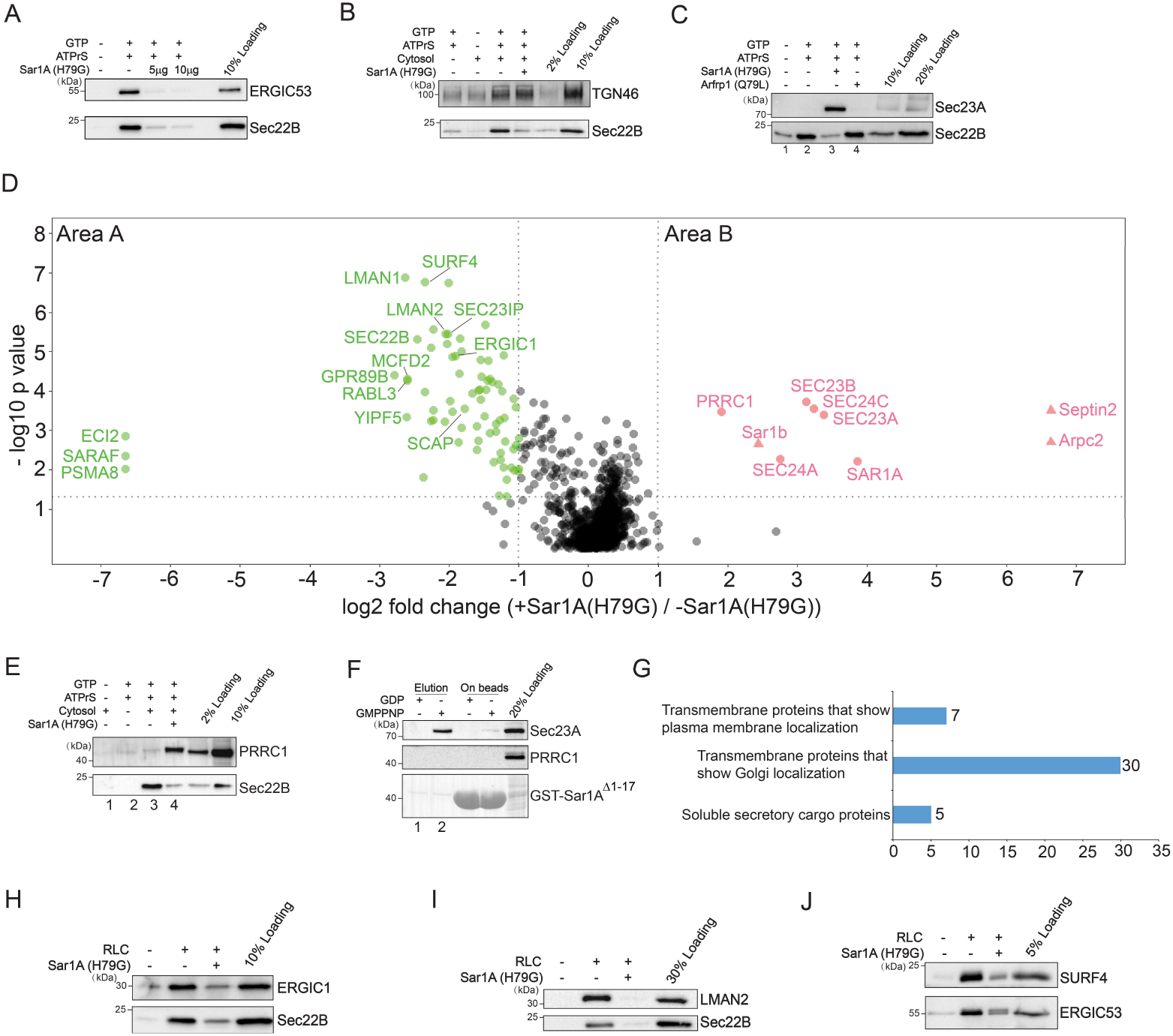
Identification of cargo proteins and cytosolic proteins that are dependent on Sar1A to be associated with transport vesicles. **A-C.** The vesicle formation assay was performed using the indicated reagents. Vesicle fractions were analyzed by immunoblot. **D.** The vesicle formation assay was performed in the presence or absence of Sar1A (H79G). The isolated vesicles in each experimental group were resuspended in RapiGest SF surfactant. The proteins in the vesicle fractions were trypsin digested and analyzed by label-free mass spectrometry. The log2 ratio of the abundance of each identified protein in the vesicles prepared in the presence of Sar1A (H79G) over that in the vesicles prepared in the absence of Sar1A (H79G) was plotted on the x axis and the −log10 p value of the difference was plotted on the y-axis. **E.** The vesicle formation assay was performed using the indicated reagents. Vesicle fractions were analyzed by immunoblot. **F.** GST-Sar1A^Δ1-17^ was loaded with GDP or GMPPNP and then incubated with rat liver cytosol. After incubation, proteins that bound to Sar1A in a nucleotide-dependent manner were eluted with EDTA. The eluted fraction and the proteins left on beads after elution was analyzed by immunoblot. **G.** Number of proteins identified in Area A of panel D categorized based on predictions from Uniprot. **H-J.** Vesicle formation assay was performed using the indicated reagents. The vesicle fraction was analyzed by immunoblot.

We propose that proteins that are significantly reduced in the presence of Sar1A(H79G) are cargo proteins associated with COPII vesicles, and that cytosolic proteins that are significantly enhanced in the presence of Sar1A(H79G) are COPII coat proteins or proteins that directly or indirectly interact with COPII coat. We therefore performed our vesicle formation assay at a large scale in the presence or absence of Sar1A(H79G). Proteins in the vesicle fractions were quantified by label-free mass spectrometry (Figure 3D). A total of 1226 proteins were identified and quantified, all of which had two or more unique peptides (FDR<0.01) and were successfully quantified in all of the three biological repeats (Table S2, sheet 1). This analysis indicates that the vast majority of proteins that are significantly enriched in vesicles generated in the presence of Sar1A(H79G) are subunits of the COPII coat (Figure 3D, Area B, Table S2, Sheet 2). However, several novel cytosolic proteins were also significantly enriched in Sar1A(H79G) condition (Figure 3D, Area B, Table S2, Sheet 2, proteins identified using the protein sequence database of *Homo sapiens* are indicated in round shapes and additional proteins identified using database of *Rattus norvegicus* are indicated in triangle shapes).

Western blot analysis confirmed the enrichment of one of these proteins, PRRC1 (Figure 3E, compare lanes 4 and 3), suggesting that PRRC1 interacts with Sar1A (H79G) on vesicle membranes. We then tested whether PRRC1 can be detected as a binding partner of Sar1A utilizing a GST-pull down approach. Purified GST-tagged human Sar1A depleted of its N-terminal amphipathic helix (GST-Sar1A^Δ1-17^) was loaded with GDP or GMPPNP and then incubated with rat liver cytosol. The concentration of rat liver cytosol in the reaction mixture was equal to that used in the vesicle formation assay. After incubation, proteins that bound to Sar1A in a GTP-dependent manner were eluted with EDTA. Western blot analysis of the eluted fraction indicated that Sec23A can be specifically detected in the eluate of GMPPNP-loaded but not GDP-loaded GST-Sar1A^Δ1-17^ immobilized on glutathione beads (Figure 3F, compare lanes 1 and 2). In contrast, PRRC1 cannot be detected in the eluate of GMPPNP-loaded GST-Sar1A^Δ1-17^ (Figure 3F). In this GST-pull down approach, the reaction was performed in the absence of lipid bilayer. In the vesicle formation assay, the lipid bilayers of the ER and the Golgi in the semi-intact cells were not disrupted and this assay was performed without detergent. Thus, the vesicle formation assay has an advantage to reveal protein-protein interactions, such as the interaction between PRRC1 and Sar1A, that take place on lipid bilayers.

Seventy-three proteins were identified with significant enrichment in the untreated group over the Sar1A(H79G) group (Figure 3D, Area A, Table S2, Sheet 4). 54 of them are predicted to be transmembrane proteins and 5 were soluble secretory cargo proteins (Figure 3G, Table S2, Sheet 5). Many of the transmembrane proteins were predicted to show plasma membrane and Golgi localization (Figure 3G, Table S2, Sheet 5). All of the SNARE proteins identified in area B (Sec22A, Sec22B, STX5, GOSR2, BET1) mediate ER-to-Golgi trafficking. Western blot analysis confirmed that three transmembrane proteins, ERGIC1, SURF4 and LMAN2 were present in the vesicle fraction and their vesicular release was significantly reduced by Sar1A (H79G) (Figure 3H-J), indicating that they are packaged in COPII vesicles. Several cytosolic proteins identified in area A including RabL3, Sec23IP, SCFD1. SCFD1 and Sec23IP have been shown to regulate ER-to-Golgi trafficking in mammalian cells (Dascher and Balch, 1996; Nogueira et al., 2014; Ong et al., 2010). The role of RabL3 in ER-to-Golgi trafficking remains to be investigated.

We then analyzed the functional role of PRRC1 in COPII-mediated ER export process. We found that knockdown of PRRC1 caused a significant enhancement of the total fluorescent level of Sec31A but not Golgin97 per cell (Figure 4A-G, quantifications in Figure 4H-I). The expression level of Sec31A in PRRC1 knockdown cells was similar to that in Mock cells (Figure 4G). These results suggest that PRRC1 downregulates membrane recruitment of Sec31A. In addition, knockdown of PRRC1 caused a delay of transport of a cargo protein, EGFR, from the ER to the Golgi in the RUSH transport system (Figure 4J and quantification in Figure 4K), indicating that PRRC1 is important for ER-to-Golgi transport of EGFR. We propose that PRRC1 regulates disassembly of COPII coat to allow the released COPII subunits to perform next round of cargo sorting on the ER membranes.

**Figure 4.**
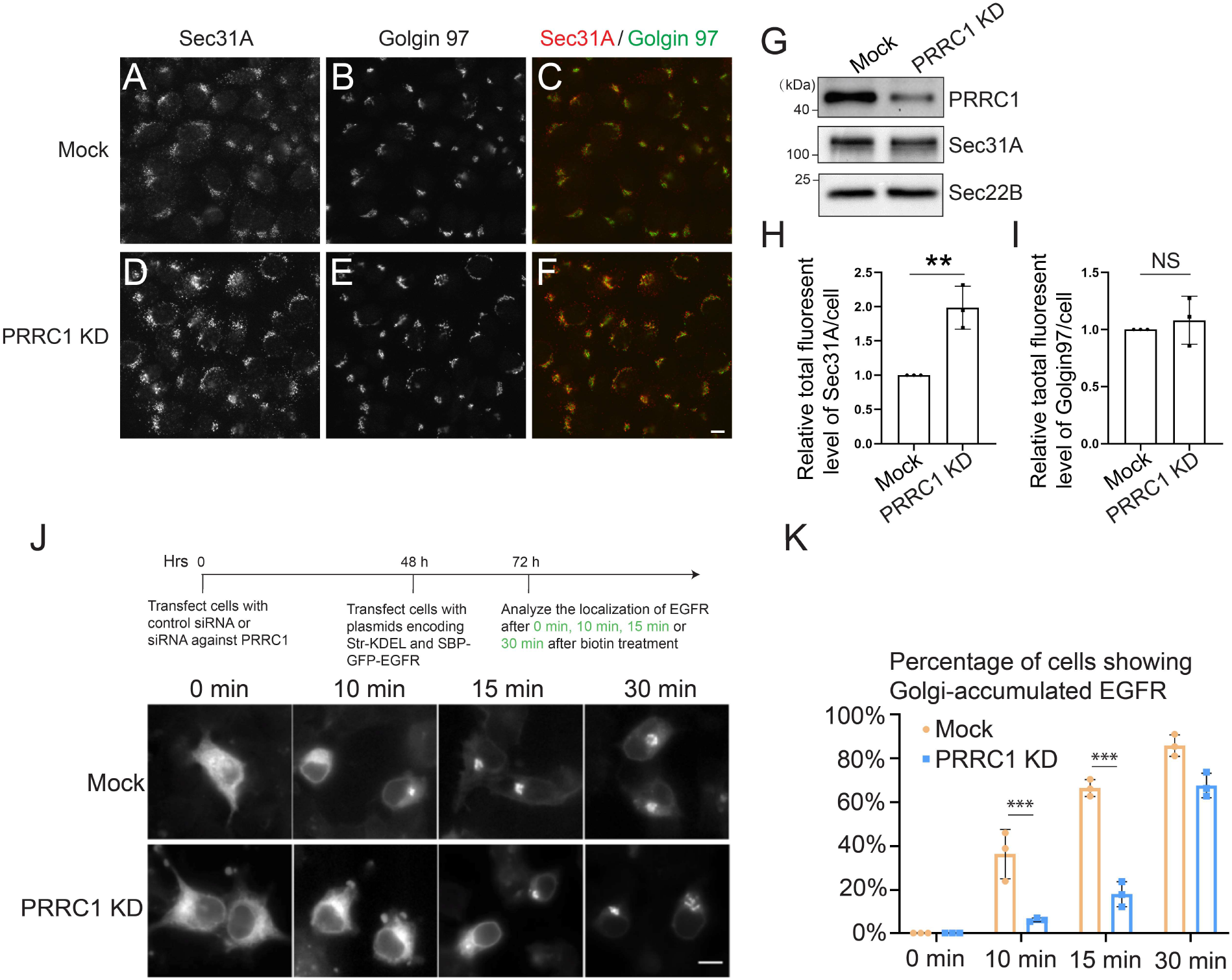
PRRC1 regulates disassembly of COPII and ER-to-Golgi transport of EGFR. **A-F.** HEK293T cells were transfected with control siRNA (A-C) or siRNA against PRRC1 (D-F). Day 3 after transfection, the localizations of Sec31A and Golgin97 were analyzed by immunofluorescence. *Scale bar*, 10 μm. **G.** HEK293T cells were transfected with control siRNA or siRNA against PRRC1. Day 3 after transfection, cells were lysed and analyzed by western blot. **H-I.** Quantifications of the total fluorescent level of Sec31A (H) and Golgin97 (I) per cell (mean ± S.D.; n = 3; >125 cells from 9 random imaging fields counted for each experiment). In each experiment, the total fluorescent level was normalized to that in Mock cells. **, p < 0.01; NS, not significant. **J.** HEK293T cells were transfected with control siRNA or siRNA against PRRC1. 24hr after transfection, cells were retransfected with plasmids encoding both EGFR^RUSH^ and Str-KDEL. On day 3 after knockdown, cells were incubated with biotin for the indicated time and the localization of EGFR^RUSH^ was analyzed by fluorescent microscope. *Scale bar*, 10 μm. **K.** Quantification of the percentage of cells showing Golgi-localized EGFR^RUSH^ in cells treated with control siRNA or siRNA against PRRC1 (mean ± S.D.; n = 3; >100 cells counted for each experiment). ***, p < 0.001.

In summary, these analyses revealed novel candidate cargo proteins that are packaged into COPII vesicles and novel cytosolic proteins that are associated with COPII-coated vesicles. Moreover, these analyses indicate that our approach is robust to reveal effector proteins that are associated with GTP-bound Arf family proteins on vesicle membranes.

### Identification of cargo proteins that depend on a specific cargo receptor for packaging into transport vesicles

Various transmembrane proteins have been implicated to function as cargo receptors to mediate packaging of soluble as well as transmembrane cargo proteins into transport vesicles. Two of these proteins, ERGIC53 and SURF4, were efficiently packaged into COPII vesicles (Figure 3J). To reveal the client repertoire of ERGIC53 and SURF4, we performed the vesicle formation assay using donor membranes prepared from genome engineered ERGIC53 knockout HEKTrex cells or SURF4 knockout HEKTrex cells (Figure 5A). Vesicles were generated from KO and WT donor membranes incubated with rat liver cytosol. Western blot analysis confirmed the absence of ERGIC53 or SURF4 in the vesicles generated from the corresponding KO donor membranes (Figure 5B-C). Proteins in the vesicle fractions were then analyzed by label free quantitative mass spectrometry. A total of 815 proteins were identified and quantified, all of which had two or more unique peptides (FDR<0.01) and were successfully quantified in all of the three biological repeats of all the experimental groups (Table S3, sheet 1). In each case, several proteins were significantly reduced in the vesicle fraction of the KO reaction compared to WT (Area A, Figure 5D-E, Table S3 sheet 2). Transmembrane proteins that are predicted to show Golgi or plasma membrane localization are highlighted in red and soluble secretory proteins highlighted in green. Additional cargo proteins were significantly reduced in the vesicle fraction in the KO vesicles compared to the WT when the threshold was changed from 0.5 to 0.6 (Area A, Figure 5D-E, Table S3 sheet 2).

**Figure 5.**
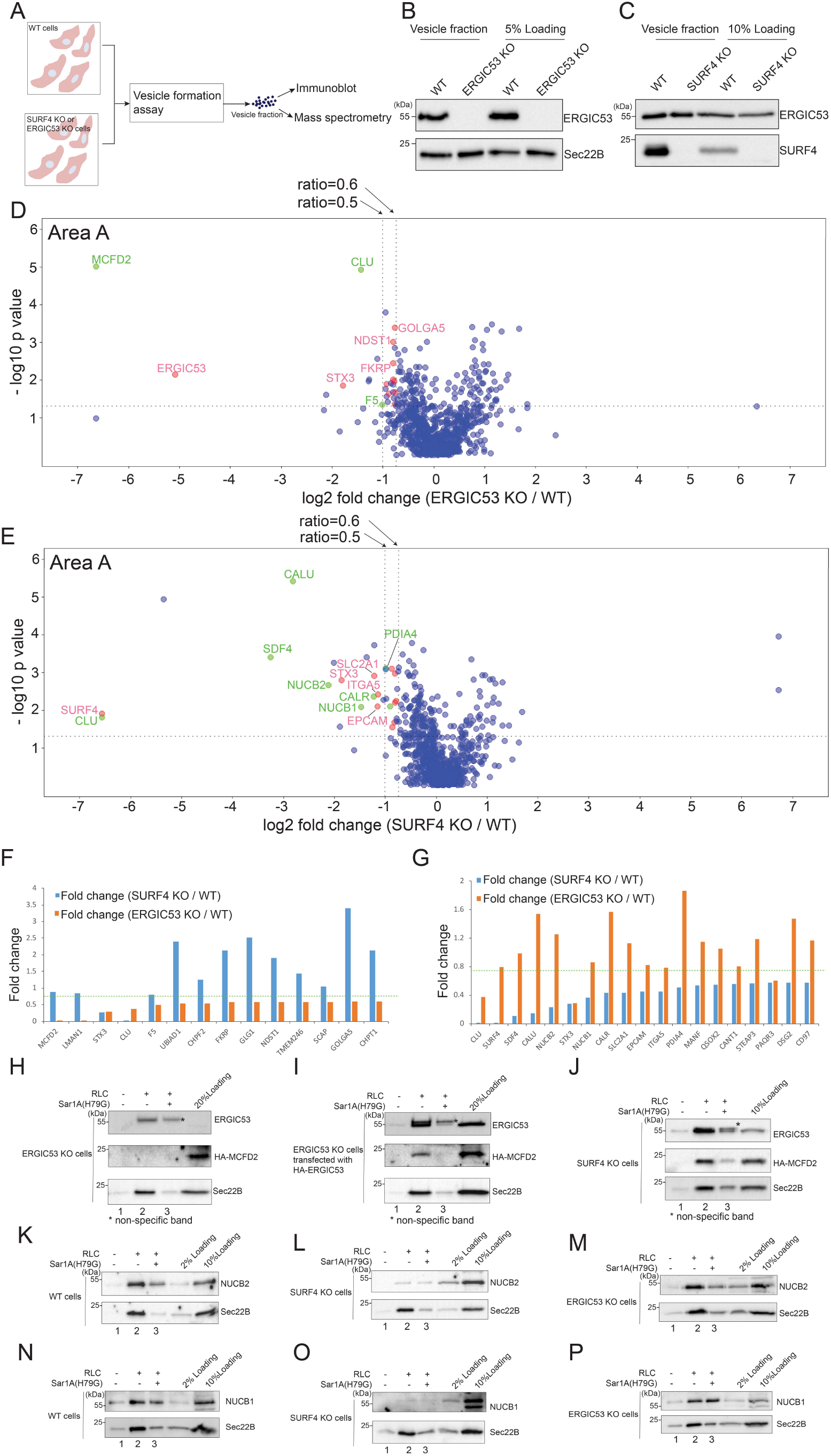
Identification of cargo proteins that depend on ERGIC53 or SURF4 for packaging into transport vesicles. **A.** Diagram demonstrating the experimental procedures. **B-C.** The vesicle formation was performed using wild type HEK293TRex cells and ERGIC53 KO HEK293TRex cells (B) or SURF4 KO HEK293TRex cells (C). The vesicle fraction was then analyzed by immunoblot. **DE.** The vesicle formation assay was performed using wild type HEK293TRex cells and ERGIC53 KO HEK293TRex cells (D) or SURF4 KO HEK293TRex cells (E). The isolated vesicles in each experimental group were resuspended in RapiGest SF surfactant. The proteins in the vesicle fractions were trypsin digested and analyzed by label-free mass spectrometry. The log2 ratio of the abundance of each identified protein in the vesicles prepared from ERGIC53 KO or SURF4 KO cells over that in the vesicles prepared in wild type cells was plotted on the x axis and the −log10 p-value of the difference was plotted on the y-axis. **F.** The fold change of the identified ERGIC53 client was compared with the fold change of the abundance of these proteins in the vesicle fraction prepared from the wt cells and SURF4 KO cells. **G.** The fold change of identified SURF4 client was compared with the fold change of the abundance of these proteins in the vesicle fraction prepared from the wt cells and ERGIC53 KO cells. **H-P.** The vesicle formation was performed using wild type HEK293TRex cells (K, N) and ERGIC53 KO HEK293TRex cells (H, M) or SURF4 KO HEK293TRex cells (J, L, O) or ERGIC53 KO cells transfected with HA-ERGIC53 (I). Vesicle fractions were then analyzed by immunoblot.

We defined ERGIC53 clients as those cargo proteins under-represented in the KO condition relative to wild-type (Fold change < 0.6; p<0.05). We then examined the abundance of these clients in vesicles made from SURF4 KO cells (Figure 5F). Most ERGIC53 clients were not similarly depleted in the SURF4 KO condition (Figure 5F, above the green line), suggesting that these cargo proteins are dependent on ERGIC53 but not SURF4 for efficient packaging into vesicles. Two such cargo proteins are known ERGIC53 interactors, MCFD2 and coagulation factor V (FV) (Figure 5F). MCFD2 forms a complex with ERGIC53 to facilitate the transport of coagulation factors V and VIII (FVIII) from the ER to the Golgi (Zhang et al., 2003; Zhang et al., 2005). Moreover, ERGIC53 is required for retention of MCFD2 in the early secretory transport pathway (Nyfeler et al., 2006). Immunoblot analysis confirmed that packaging of HA-tagged MCFD2 into vesicles was abrogated in ERGIC53 KO cells (Figure 5H, lane 2) but not in SURF4 KO cells (Figure 5J, lane 2). Exogenously expressing ERGIC53 in ERGIC53 KO cells rescued packaging of MCFD2 into transport vesicles, and adding Sar1A(H79G) blocked this rescue (Figure 5I, lanes 2 and 3).

Using similar criteria, we identified several SURF4 clients that were unaffected by loss of ERGIC53 (Figure 5G). Two of the top hits, NUCB1 and NUCB2, were verified to be packaged into vesicles and the efficiency was reduced by Sar1A(H79G) (Figure 5K and N, compare lanes 2 and 3). Efficiency of NUCB1 and NUCB2 packaging was greatly reduced in SURF4 KO cells (Figure 5L and O, lane 2), whereas vesicular release of these two cargo proteins was not affected in ERGIC53 KO cells (Figure 5M and P, lane 2). Altogether, our analyses revealed specific transmembrane and soluble cargo proteins that depend on SURF4 or ERGIC53 to be packaged into transport vesicles. These analyses indicate that our method is a robust approach to reveal the clients of a specific transmembrane cargo receptor.

### Identification of proteins that are specifically present in vesicles enriched with a specific cargo

Transport vesicles with specific cargo proteins should also recruit additional proteins such as Rabs, motor proteins and SNAREs to ensure vesicle targeting. To investigate this, we sought to immunoisolate vesicles enriched with a specific cargo protein, either EGFR or Frizzled6, and then perform mass spectrometry analysis to reveal other proteins that are present in the immune-isolated vesicles. We have previously characterized reconstituted release of EGFR and Frizzled6 into transport vesicles (Ma et al., 2018; Mao et al., 2019). We thus performed this assay at a large scale using HEK293T cells transfected with plasmids encoding FLAG-tagged Frizzled6 or EGFR (the Frizzled6-FLAG group or the EGFR-FLAG group) (Figure 6A). The vesicle fractions were then incubated with agarose beads conjugated with mouse anti-FLAG antibodies. Vesicles that bound to beads were eluted using FLAG peptides and analyzed by negative stain electron microscope and immunoblot. We used untransfected HEK293T cells as a negative control. We detected small membrane-bound structures in the immune-isolated fraction from the Frizzled6-FLAG and EGFR-FLAG groups but not in the control group (Figure 6B-D). Interestingly, the average diameter of the vesicles from the Frizzled6-FLAG group were significantly larger than that from the EGFR-FLAG group (Figure 6E, 69.2 ±17.0 nm vs. 57.2 ± 11.8 nm).

**Figure 6.**
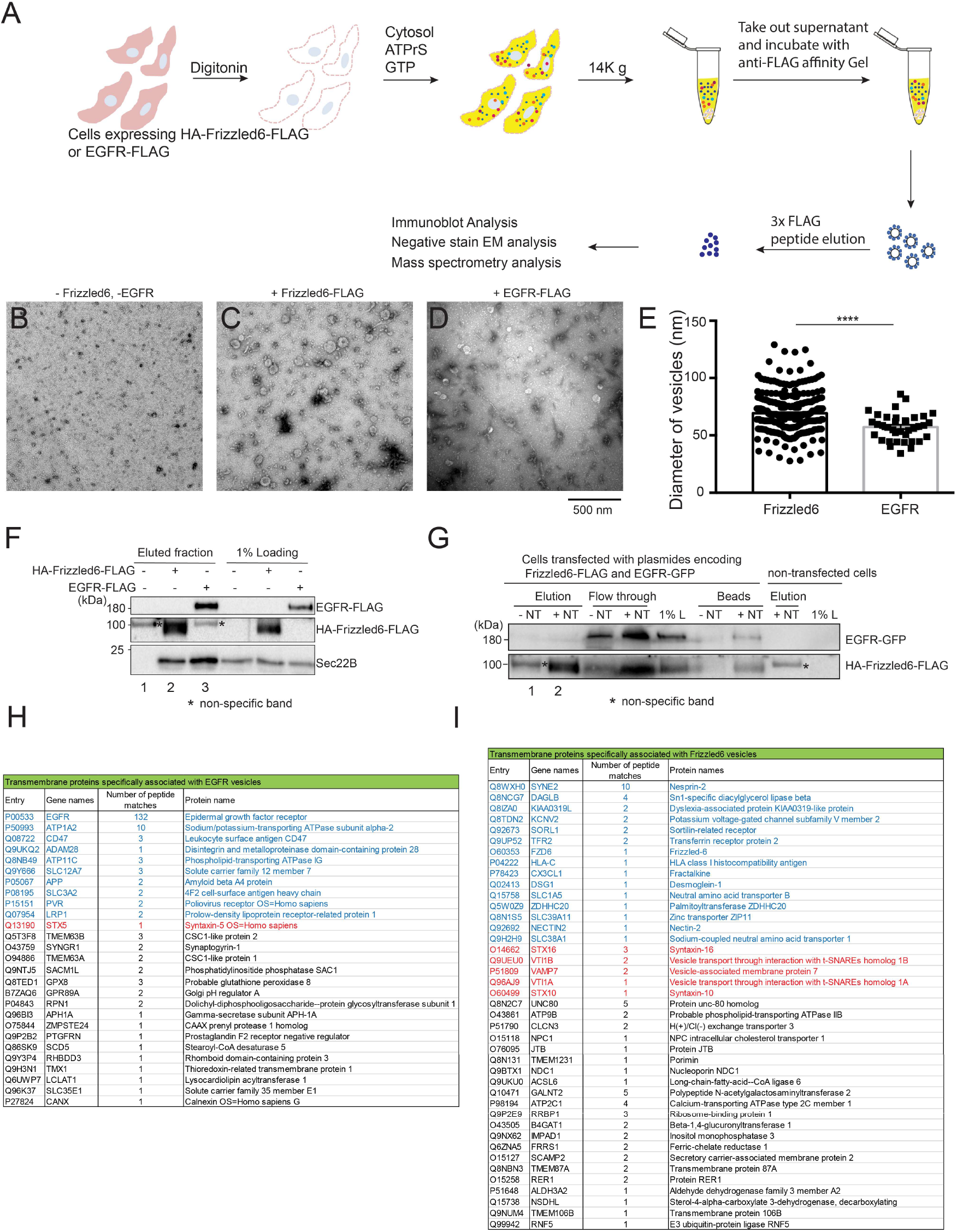
Identification of proteins that are co-packaged with Frzzled6 or EGFR in transport vesicles. **A.** A diagram demonstrating the approach to immunoisolate vesicles enriched with EGFR and Frizzled6. **B-D.** Negative stain EM analysis of the immunoisolated vesicles from untranfected cells (B) or from cells transfected with plasmids encoding Frizzled6-FLAG (C) or EGFR-FLAG (D). *Scale bar*, 500 nm. **E.** Quantification analysis of the average diameter of the donut-shaped structures observed in the immunoisolated vesicles enriched with EGFR-FLAG or Frizzled6-FLAG (mean ± SD). ****, p<0.0001. **F.** Immunoblot analysis of the immunoisolated vesicles from untransfected cells or cells transfected with Frizzled6-FLAG or EGFR-FLAG. **G.** Immunoblot analysis of the immunoisolated vesicles from non-transfected cells or from cells co-transfected with the indicated plasmids. **H.** Table showing transmembrane proteins that are identified to be co-packaged with EGFR-FLAG in vesicles. **I.** Table showing transmembrane proteins that are identified to be co-packaged with Frizzled6-FLAG in vesicles. Transmembrane proteins that are predicted to show plasma membrane localizations were highlighted in blue and SNARE proteins were highlighted in red.

Immunoblot analysis indicated that vesicles containing EGFR-FLAG or Frizzled6-FLAG were successfully immunoisolated, and that these vesicles also contained the SNARE protein, Sec22B (Figure 6F). We next performed this assay using cells co-transfected with EGFR-GFP and Frizzled6-FLAG. Immunoisolated vesicles containing Frizzled6-FLAG lacked EGFR-GFP (Figure 6G, lane 2), suggesting that EGFR and Frizzled6 are packaged in separate vesicles. Proteins in the immunoisolated vesicle fractions from untransfected control cells, or cells expressing either Frizzled6-FLAG or EGFR-FLAG were then analyzed by mass spectrometry. A variety of proteins were specifically present in vesicles enriched with ERFR-FLAG or Frizzled6-FLAG, including 27 transmembrane proteins present in EGFR vesicles (Figure 6H), 10 of which were predicted by Uniprot to show plasma membrane localizations (Figure 6H, highlighted in Blue). An additional 41 transmembrane proteins were specifically present in Frizzled6 vesicles (Figure 6I), and 14 of these were predicted plasma membrane proteins (Figure 6I, highlighted in Blue). We hypothesize that these plasma-membrane localized transmembrane proteins correspond to cargo proteins that are specifically packaged together with EGFR or Frizzled6. Interestingly, we identified several SNAREs that were specifically present in Frizzled6 vesicles but not in EGFR vesicles (Figure 6I, highlighted in red), suggesting that vesicles enriched with specific cargo proteins also contain specific SNAREs and other cargo molecules. Enrichment of specific proteins together into transport vesicles may be important for the efficient cargo sorting and the downstream vesicular targeting process.

### Vti1B is packaged together with Frizzled6 and promotes release of Frizzled6 into transport vesicles

One of the SNARE proteins identified in Frizzled6 vesicles is Vti1B which directly interacts with a TGN/endosome-located cargo adaptor, epsinR (Chidambaram et al., 2004; Chidambaram et al., 2008; Hirst et al., 2004; Miller et al., 2007; Wang et al., 2011). Immunoblot analysis verified that Vti1B is present in Frizzled6 vesicles (Figure 7A, lane 2). EpsinR contains an N-terminal ENTH domain followed by an unfolded region. The ENTH domain directly binds various SNAREs, including the Habc domain of Vti1B (Chidambaram et al., 2004; Chidambaram et al., 2008; Hirst et al., 2004; Miller et al., 2007; Wang et al., 2011). The C-terminal unfolded domain of epsinR interacts with the polybasic motif on Frizzled6 to package Frizzled6 into transport vesicles (Ma et al., 2018). We immunoisolated FLAG-tagged C-terminal unfolded domain of epsinR (epsinR^ΔENTH^-FLAG) and full length epsinR (epsinR^FL^-FLAG) from HEK293T cells. Using *in vitro* binding assays, we found that recombinant purified GST-tagged Frizzled6 C-terminal domain (GST-Frizzled6^497-567^) binds epsinR^ΔENTH^-FLAG with a higher affinity than epsinR^FL^-FLAG (Figure 7B, compare lanes 1 and 2). Interestingly, when the *in vitro* binding assay was performed in the presence of purified recombinant His-tagged Vti1B Habc domain (His-Vti1B^Habc^), the interaction between GST-Frizzled6^497-567^ and full length epsinR was enhanced (Figure 7B, compare lanes 1 and 3). Increasing the concentration of His-Vti1B^Habc^ increased the affinity between Frizzled6 and epsinR^FL^-FLAG (Figure 7C, lanes 1-4). In contrast, His-Vti1B^Habc^ did not enhance the interaction between GST-Frizzled6^497-567^ and the C-terminal unfolded domain of epsinR (Figure 7B, compare lanes 2 and 4). These results suggest that Vti1B promotes the interaction between epsinR and Frizzled6. To test whether this process enhances the efficiency of packaging of Frizzled6 into vesicles, we performed the vesicle formation assay in the presence of His-Vti1B^Habc^ We found that purified His-Vti1B^Habc^ enhanced the efficiency of packaging of Frizzled6 into vesicles (Figure 7D, compare lanes 2 and 3). Increase the concentration of His-Vti1B^Habc^ in the vesicle formation assay increased the efficiency of vesicular release of Frizzled6 (Figure 7E, compare lanes 2-5).

**Figure 7.**
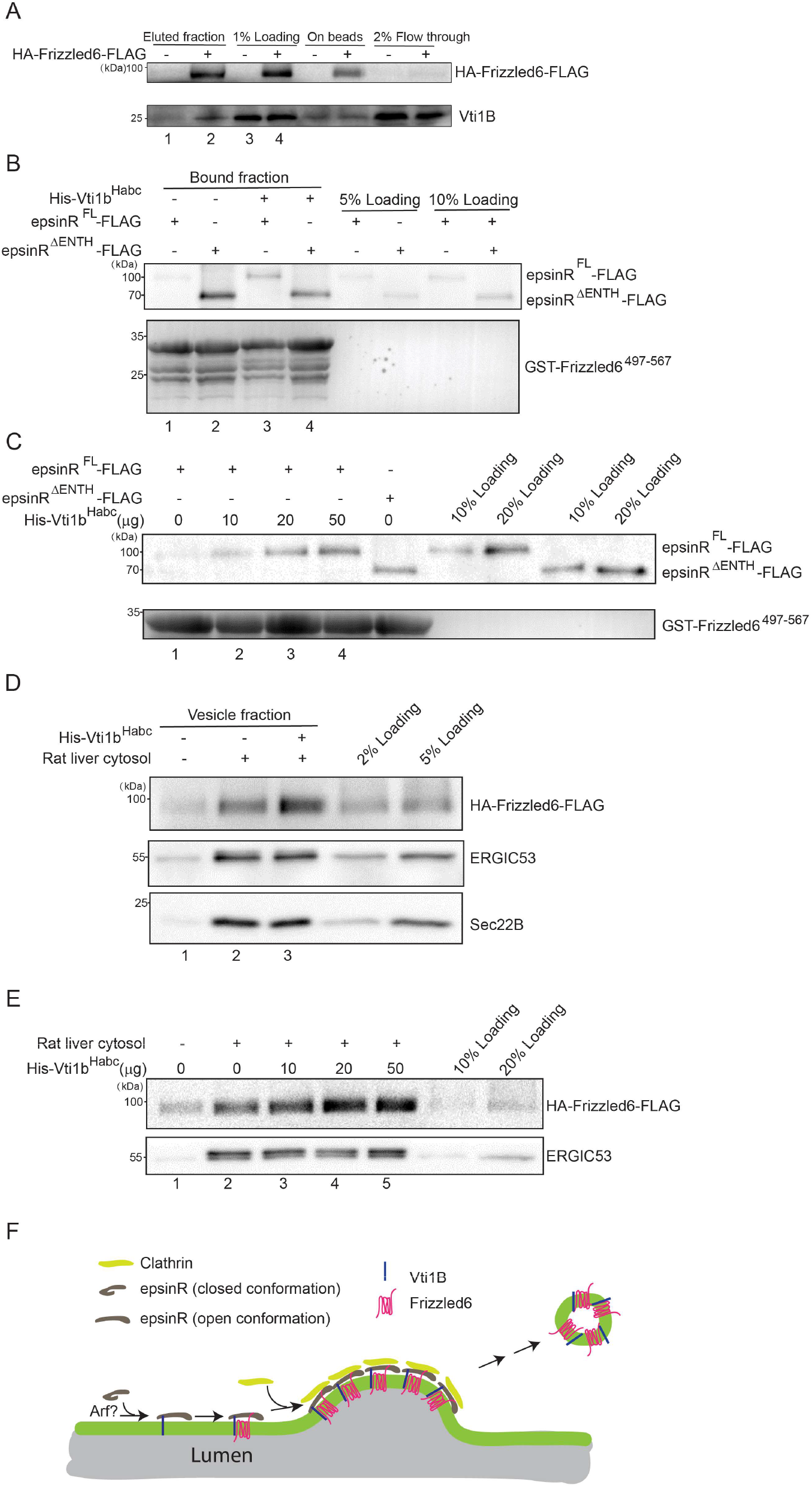
Vti1B is present in Frizzled6 vesicles, promotes binding of Frizzled6 to epsinR, and enhances the efficiency of release of Frizzled6 into vesicles. **A.** Immunoblot analysis of the immunoisolated vesicles from untranfected cells or cells transfected with Frizzled6-FLAG. **B-C.** GST-pull down analysis by incubating purified GST-Frizzled6^497-567^ with the indicated immunoisolated epsinR constructs in the presence or absence of purified His-Vti1B^Habc^ **D-E.** HEK293T cells were transfected with plasmids encoding HA-Frizzled6-FLAG. Day 1 after transfection, the vesicle formation was performed using the indicated reagents. **F.** A model demonstrate that binding of Vti1B to the ENTH domain of epsinR causes a conformational change of epsinR, allowing epsinR to bind Frizzled6 so that Vti1B and Frizzled6 are packaged together in transport vesicles. Each result is a representative figure from three biological repeats.

Next, we analyzed whether Vti1B is important for cell surface delivery of Frizzled6 in HeLa cells through a temperature shift approach (Ma et al., 2018). In this approach, we incubated HeLa cells at 20 °C in the presence of cycloheximide to accumulate newly-synthesized HA-Frizzled6 in the TGN. A surface-labeling experiment was performed to label the HA tag localized on the extracellular domain of Frizzled6. After a 20 °C incubation, HA-Frizzled6 was accumulated at the juxtanuclear Golgi area and the majority of cells showed no detectable surface-localized Frizzled6 (Ma et al., 2018). After 32 °C incubation, Frizzled6 in most control siRNA-treated cells showed a detectable surface-localized pattern (Ma et al., 2018). Utilizing this approach, we analyzed the efficiency of TGN-to-cell surface delivery of Frizzled6 in control cells or cells transfected with siRNA against Vti1B which efficiently reduced the expression level of Vti1B (Figure S2G). We quantified the ratio of the fluorescent level of surface-localized Frizzled6 over the fluorescent level of total Frizzled6 after 32 °C incubation and this ratio was used as an indicator of the efficiency of transport. We found that knockdown of Vti1B caused a significant reduction of Surface/Total ratio of Frizzled6 (Figure S2A-F, quantification in Figure S2H), suggesting that Vti1B regulates delivery of Frizzled6 to the cell surface.

These results indicate that epsinR has a closed conformation in which the ENTH domain blocked the Frizzled6-binding site on the epsinR C-terminal domain (Figure 7F). Binding of Vti1B to the ENTH domain of epsinR causes a conformational change of epsinR. This process exposes the cargo binding site thereby promoting the interaction between epsinR and Frizzled6 to capture Frizzled6 into vesicles (Figure 7F). We hypothesize that this mechanism allows a specific cargo protein, Frizzled6, to be copackaged together with a specific SNARE, Vti1B, to ensure the downstream vesicle targeting process.

## Discussion

Protein sorting is an essential step to ensure a correct delivery of specific cargo proteins to their specific destination. Although significant progresses have been achieved, the specificity of the protein sorting process mediated by a specific cellular factor remains to be further investigated. In addition, vesicular trafficking is regulated by a variety of cytosolic proteins which are recruited to membranes to regulate important processes such as capturing cargo proteins, regulating vesicle coat assembly or linking vesicles to cytoskeleton. Revealing these cytosolic factors will provide important insight into the vesicular trafficking process. Our study demonstrates that the vesicle formation assay in combination with quantitative mass spectrometry analysis is a robust and powerful tool to investigate these aspects.

GTP-binding proteins, including Arf family proteins and Rab family proteins, play critical roles in mediating membrane recruitment of cytosolic factors to regulate cargo sorting and vesicle formation (Christoforidis and Zerial, 2000; Guo et al., 2013; Jin et al., 2010). Affinity chromatography is a traditional approach to identify these cytosolic factors. Here, we utilized a novel approach that preserves membranes in the interaction. A flotation step efficiently removes the majority of cytosolic proteins that are not tightly associated with vesicles. Our analysis indicates that our approach can reveal novel protein-protein interactions that take place on lipid bilayers. Through this approach, we identified several cytosolic factors that are associated with vesicle membranes in a GTP-dependent manner or that interact with GTP-bound Sar1A on vesicle membranes. These cytosolic proteins may function as cargo adaptors or may associate with vesicle coats to regulate cargo sorting. In addition to Sar1A, we identified several further Arf and Rab proteins whose abundance was significantly increased in the vesicle fraction produced in the presence of GMPPNP. It would be interesting to utilize our approach to reveal the cytosolic proteins and cargo proteins that depend on these proteins to be associated with transport vesicles (Figure 8A).

**Figure 8.**
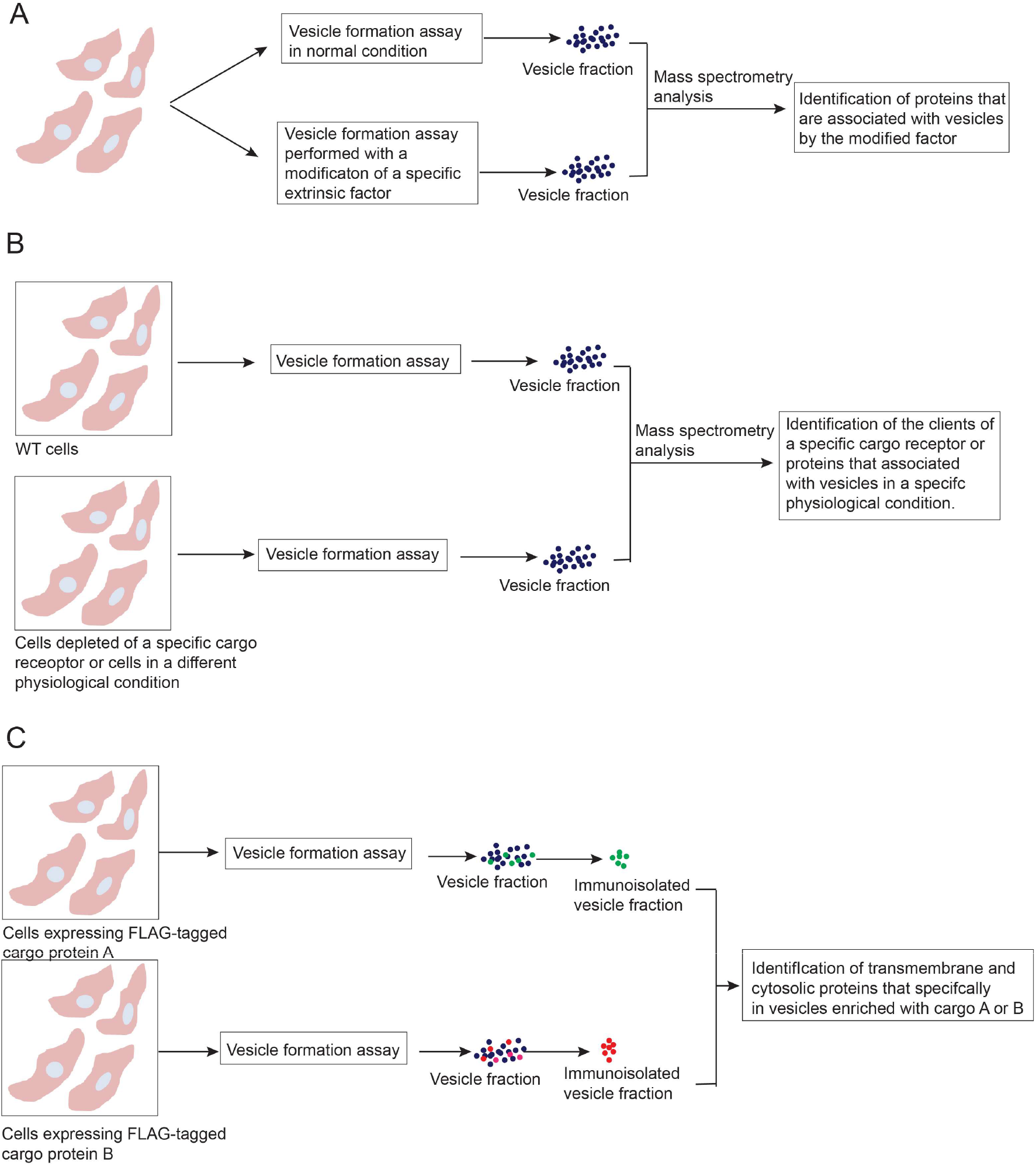
The applications of the vesicle formation assay to analyze the protein sorting process in the secretory pathway. **A.** By modification of a specific extrinsic factor in the vesicle budding reaction, this approach can reveal cytosolic proteins or cargo proteins that are associated with vesicles by the modified factor to regulate vesicular trafficking. **B.** By depleting a specific cargo receptor or using cells under different physiological conditions, this approach can reveal the clients of a specific cargo receptor or reveal novel proteins that associated with vesicles to regulate a specific physiological process. **C.** By immunoisolate vesicles enriched with a specific cargo protein, this approach can provide a better understanding of the protein composition of transport vesicles enriched with a specific cargo to provide insights into the cargo sorting and vesicle targeting process.

Two of these identified cytosolic proteins, FAM84B and PRRC1, were shown to regulate ER-to-Golgi transport of newly synthesized EGFR. FAM84B contains a LRAT (lecithin: retinal acyltransferase) domain. This domain is present in the H-Ras-like suppressor (HRASLS) family. The expression of FAM84B is upregulated during prostate cancer progression and in preclinical and Esophageal squamous cell carcinoma tumors (Cheng et al., 2016; Wong et al., 2017). FAM84B is shown to promote prostate tumorigenesis (Jiang et al., 2019). PRRC1 is predicted to have protein kinase A regulatory subunit binding activity. Our results indicate that PRRC1 localizes to the endoplasmic reticulum and downregulates membrane recruitment of the outer COPII subunit, Sec31A. PRRC1 contains a proline-rich domain. The proline-rich region of Sec31 interacts with Sec23 (Shaywitz et al., 1997; Shugrue et al., 1999), suggesting that PRRC1 may directly interact with Sec23 to perform its function.

Our study demonstrates that the vesicle formation assay is powerful to reveal specific cargo proteins that depend on distinct cargo receptors to be efficiently packaged into transport vesicles (Figure 8B). This analysis will not only provide important information on the specificity of protein sorting but also reveal insight into the functional role of a specific cargo receptor. Another application of our approach is to analyze the protein composition of the vesicles produced from cells under different physiological conditions such as starvation (Figure 8B). This could provide important insight into how vesicles contribute to establish and maintain a specific physiological condition. A caveat of our assay is that this assay relies on identification of cargo proteins that are actively produced by cells. It would be interesting to perform this assay in cell lines that highly secret a variety of cellular factors to identify cargo proteins.

Transport vesicles should contain both cargo proteins and proteins involved in vesicle movement and targeting. Our analysis indicates that the vesicle formation assay is robust to analyze the protein composition of transport vesicles enriched with a specific cargo (Figure 8C), which will provide insight into how the cargo sorting machinery is coupled to the vesicle targeting machinery. A specific transport vesicles enriched with TGN46 has been immunoisolated and characterized (Wakana et al., 2012). These vesicles contain other secretory cargoes and plasma membrane localized cargoes but not vesicular stomatitis virus (VSV)-G protein and collagen I (Wakana et al., 2012). These vesicles also contain myosin II, Rab6a, Rab8a, and synaptotagmin II but whether these proteins are also enriched in vesicles enriched with other cargo proteins are not clear (Wakana et al., 2012). In this study, we demonstrated that vesicles enriched with a plasma-membrane localized signaling receptor, Frizzled6, are specifically enriched with specific SNARE proteins that may regulate vesicle targeting. We also provide mechanistic analysis to provide insights into how a SNARE protein, Vti1B, is co-enriched with Frizzled6 in vesicles (Figure 7F). An important application of our approach is to investigate the protein composition of vesicles enriched with cargo proteins that are delivered to specific destination such as apical plasma membranes or basolateral plasma membranes. This analysis will provide important information to understand how vesicles are targeted to their specific destination.

## Materials and Methods

### Cell lines, antibodies and plasmids

HEK293T and HeLa cell lines were kindly provided by the University of California-Berkeley Cell Culture Facility. HEK 293Trex, HEK 293Trex ERGIC53 KO and HEK 293Trex SURF4 KO cell lines were kindly provide by Prof. Liz Miller’s lab (MRC Laboratory of Molecular Biology, UK). All cell lines were tested negative for mycoplasma contamination. HeLa and HEK293T cells were maintained in Gibco Dulbecco’s Modified Eagle Medium (DMEM) containing 10% fetal bovine serum (FBS) and 1% penicillin streptomycin mix. HEK 293Trex, HEK 293Trex ERGIC53 KO and HEK 293Trex SURF4 KO cell lines were maintained in Gibco DMEM containing 5 μg/ml blasticidin and 10% FBS. For temperature shift experiments, HEK293T, HEK 293Trex, HEK 293Trex ERGIC53 KO or HEK 293Trex SURF4 KO cells were incubated in Opti-MEM (Invitrogen, NY) containing 10% FBS at 15°C for 2.5 hrs to accumulate cargo proteins at the ER.

For CRISPR experiments, sgRNA sequences ligated into pX458 (pSpCas9 BB-2A-GFP) plasmids were purchased from GenScript. Transfections were performed with TransitIT-293 (Mirus Bio) per manufacturer’s instructions. Clonal cell lines were derived by diluting cell suspensions to a single cell per well and expanding individual wells. Genotyping of clonal cell lines was performed by Sanger sequencing of target site PCR amplicons of genomic DNA isolated by Puregene kit (Quiagene). sgRNAs were as follows: SURF4: 5’AGTCGCGCTGCTCGCTCCAC3’ targeting exon 1; ERGIC53: 5’TGACGGGGCTAGTCAAGCTT3’ targeting exon 4.

Plasmids encoding His-tagged human Arf1, Arfrp1, Sar1A in pET28a vector and HA-tagged mouse Vangl2 in pCS2 vector were kindly provided by the Schekman lab (University of California, Berkeley). Plasmids encoding HA-Frizzled6-FLAG was generated as described (Ma et al., 2018). Plasmids encoding EGFR-FLAG and FAM84B-FLAG were generated by inserting human EGFR or human FAM84B into p3xFLAG-CMV-14 vector. Plasmids encoding HA-tagged human MCFD2 and human PRRC1 in pcDNA3.1(+) vector were synthesized by Beijing Genomics Institution (BGI). Plasmids encoding Str-KDEL_SBP-EGFP-EGFR were generated by replacing the DNA fragment encoding E-cadherin within the plasmid Str-KDEL_SBP-EGFP-Ecadherin (Addgene, Plasmid #65286) with a DNA fragment encoding human EGFR (amino acids 31-1210).

The polycolonal rabbit antibodies against Sec22B, ERGIC53, Sec23A/B were gifts from Dr. Randy Schekman’s lab (UC Berkeley). Rabbit anti-AP1μ1 antibodies were gifts from Dr. Jim Hurley’s lab (UC Berkeley). Rabbit anti-SURF4 antibodies were gifts from Dr. Xiaowei Chen’s lab (Peking University). The commercial antibodies were: mouse anti-AP1γ1 (BD Bioscience, number 610385, RRID: AB_397768); Mouse anti-His (Qiagen, number 34660); Mouse anti-β-actin (Proteintech, number 60008-1-Ig); Rabbit anti-LMAN2 (Abcam, number ab124146); Rabbit anti-HA (Cell signaling technology, number 3724S); Sheep anti-TGN46 (AbD Serotec, number AHP500G); Rabbit anti-PRRC1 (Bethyl Laboratories, number A305-783A-T); Rabbit anti-FAM84B (Proteintech, number 18421-1-AP); Rabbit anti-NUCB1 (Abcam, number ab206697); Rabbit anti-NUCB2 (Abcam, number ab229683); Rabbit anti-TMED10 (Proteintech, number 15199-1-AP).

Small interfering RNAs (siRNAs) were purchased from RUI BO (Guangzhou, China). Cells were harvested after 48 hrs or 72 hrs transfection.

The target sequence against PRRC1 is GACAAAACATTCAGTAGAA. The target sequence against FAM84B is GCAACCAGGTGGAGAAATT. The target sequence against TMED10 is GTGAGGAGATTCACAAGGA.

### Transfection and Immunofluorescence Staining

DNA constructs were transfected into HeLa, HEK293T, HEK 293Trex, HEK 293Trex ERGIC53 KO or HEK 293Trex SURF4 KO cells using lipofectamine 2000 (Invitrogen) or polyethyleneimine (PEI). Transfection of siRNA into HEK293T cells or HeLa cells was also performed using lipofectamine 2000 as described in the protocol provided by Invitrogen. The final working concentration of each siRNA is 40 nM.

Immunofluorescence was performed as described (Ma et al., 2018). HEK293T cells or HeLa cells were plated at 13 mm coverslip. Cells were fixed by incubating in 4% paraformaldehyde (PFA) in PBS at room temperature for 20 min. After fixation, cells were incubated with blocking buffer (2.5% FBS, 0.1%Triton X-100 and 0.2 M Glycine in PBS) for 30 min at room temperature. Cells were then incubated in primary antibody in blocking buffer at room temperature for 1 hr and washed 3 times with PBS, followed by incubating with indicated secondary antibodies for 30 min in blocking buffer at room temperature. Cells were washed 3 times with PBS before being mounted onto slides with ProLong antifade mountant (Vector Laboratories). Surface labeling of Frizzled6 was performed as described (Ma et al., 2018). Images were acquired with a Zeiss Axioobserver Z1 microscope system.

Quantifications of the total fluorescence of Sec31A, Gogin97, surface localized HA-Frizzled6 and total HA-Frizzled6 were performed as described using Image J (Guo and Linstedt, 2006). For each experiment, a fixed threshold was manually selected that covers most of the signal on the original gray-scale images and applied to all images. Individual cells were then selected with the free-hand tool and the total above-threshold fluorescence was determined using the measure function.

### *In vitro* vesicle formation assay

In vitro vesicle formation assay was performed as described (Ma et al., 2018). HEK293T, HEK 293Trex, HEK 293Trex ERGIC53 KO or HEK 293Trex SURF4 KO cells were permeabilized in ice cold KOAc buffer (110 mM potassium acetate, 2.5 mM magnesium acetate, 20 mM Hepes, pH 7.2) containing 40 mg/mL digitonin on ice for 5 min. The permeabilized cells were collected by centrifugation at 300 g for 3 min at 4 °C, washed with KOAc buffer and resuspended in KOAc buffer. The semi-intact cells were then incubated at 30 °C with 2mg/ml rat liver cytosol (RLC), 200 μM GTP or GMPPNP (Wako, number SAH3766) and an ATP regeneration system consisting of 4mM creatine phosphate (Roche), 0.02 mg/ml of creatine phosphokinase (Roche), and 100 μM ATP (Sigma). After a 1 hr incubation, the reaction mixture was centrifuged at 14000 g at 4 °C for 20 min to remove the ER and Golgi membranes, nucleus as well as other cell debris from the reaction mixture. The supernatant fraction containing the released vesicles was resuspend in 35% Opti-Prep and overlaid with 30% Opti-Prep. 50 μL KOAc buffer was added on the top of the step gradient of Opti-Prep. The Opti-Prep gradients were then centrifuged at 100000 g in a TLS55 (Beckman) or S55S (Hitachi) rotor at 4°C for 1.5 hr. After centrifugation, the top fraction (the vesicle fraction) was collected and was analyzed by negative stain electron microscope, Coomassie staining, western blot or mass spectrometry.

To immunoisolate vesicles enriched with Frizzled6 or EGFR, the vesicle formation assays were performed using Frizzled6-FLAG or EGFR-FLAG overexpressed HEK293T cells. After 14000 g centrifugation, the supernatant fraction containing the released vesicles was incubated with anti-FLAG M2 Affinity Gel (Sigma-Aldrich, Cat number: A2220) and 1 mM DTT in the presence of proteinase inhibitors at 4 °C with rotation overnight. The gel was then washed by KOAc buffer for 6 times and eluted by KOAc buffer containing 0.6 mg/ml 3x FLAG peptides. The eluted fraction was analyzed by immunoblot and Coomassie Blue staining.

### Negative stain electron microscopy analysis

Negative stain electron microscopy analysis was performed essentially as described previously (Tse et al., 2004). Briefly, 10 μl of the vesicle fraction was air dried onto the Carbon-coated 400-mesh copper grids, which were pre-glow discharged using PELCO easiGlowTM Glow Discharge Cleaning System (Ted Pella, Inc., Redding, CA). The copper grids containing vesicles were then negatively stained with 5% uranyl acetate for 2 min and further dried before observation. The vesicle samples were examined using a Hitachi H-7650 transmission electron microscope with a CCD camera operating at 80 kV (Hitachi High-Technologies Corporation, Japan).

### Purification of His tagged proteins from E. coli

Cells expressing His-tagged constructs were grown to the optical density at 600 nm (OD600) at 0.6-0.8 in lysogeny broth (LB) and induced with 0.5mM isopropyl β-D-1-thiogalactopyranoside (IPTG) at 25 °C for 4 hr for the protein expression. The cells were lysed with lysis buffer containing 10mM imidazole and 1mg/ml lysozyme in 2x PBS buffer (274mM NaCl, 5.4mM KCl, 20mM Na2HPO4, 3.6mM KH2PO4, pH7.4), followed by the sonication. The lysate was centrifuged at 100000 g for 10 min to remove cell debris and then the supernatant was incubated with HisPur™ Ni-NTA Resin overnight. The protein was eluted by elution buffer containing 250 mM imidazole, protease inhibitors and 1mM Dithiothreitol (DTT) in KOAc buffer. The eluted protein was dialyzed against KOAc buffer.

### Nucleotide loading and GST pull down

Purification of GST-tagged protein was performed as descripted (Guo et al., 2013). The nucleotide loading and GST pull down experiment was performed as descripted (Christoforidis and Zerial, 2000; Guo et al., 2013). Briefly, 10 μl glutathione beads bearing around 10 μg GST-Sar1A^Δ1-17^ were washed 3 times with 200 μl Nucleotide Exchange (NE) buffer (20 mM HEPES, pH 7.4, 100 mM KCl, 2mM EDTA, 1mM MgCl2, 1 mM dithiothreitol) then the beads were incubated with 150 μl NE buffer containing 500 μM GDP or GMPPNP at 37 °C for 90 min. After incubation, the beads were washed 3 times with 200 μl Nucleotide Stabilization (NS) buffer (20 mM HEPES, pH 7.4, 100 mM KCl, 5mM MgCl2, 1 mM dithiothreitol) then incubated with 150 μl NS buffer containing 500 μM GMPPNP or GDP at 37 °C for 45 min. After incubation, the beads were incubated with 2 mg/ml rat liver cytosol containing 100 μM GDP or GMPPNP at 4 °C for 2hrs. The beads were then washed six times with 500 μl NS buffer and one time with 500 μl NS buffer without magnesium and then the bound proteins were eluted with elution buffer (20mM Hepes, 1.5M KCl, 0.5% Triton-100, 20mM EDTA) and analyzed by immunoblotting. All of the incubations were performed in the presence of proteinase inhibitors.

### Retention Using Selective Hook (RUSH) assay

HEK293T cells were transfected with plasmids encoding Str-KDEL_SBP-EGFP for 24 hrs. To release the SBP-EGFP-EGFR from the ER, cells were treated with 100 ng/μl cycloheximide (Sigma-Aldrich) and 40 μM D-biotin (Sigma-Aldrich) for the indicated time. Cells were then washed with PBS, fixed with 4% PFA, washed with PBS and mounted onto slides with ProLong antifade mountant (Vector Laboratories).

### Labeled-free quantitative mass spectrometry

0.1% RapiGest SF Surfactant was dissolved in 50 mM triethylammonium bicarbonate (TEAB) (Sigma-Aldrich, number T7408). Vesicle samples were resuspended in 0.1% RapiGest by vortexing. The same volume of 8M urea dissolved in 50 mM TEAB buffer was added into the vesicle sample. The sample was then reduced with 10mM TCEP at 37°C for 1 hr and alkylated with 20mM Indole-3-acetic acid (IAA) at room temperature in the dark for 30 min, followed by the digestion with sequencing grade modified trypsin (Promega, number V511A) at 37 °C for 20 hr. To stop the trypsin digestion and to remove the surfactant, the pH of vesicle samples was adjusted to 2.5-3.0 by adding 10% trifluoroacetic acid (TFA). The degraded surfactant was removed by centrifugation. Samples were dried by speed vacuum. Next, samples were desalted with pierce C18 spin column. Subsequently, the samples were dried again to be analyzed by the mass spectrometry.

### Mass spectrometry and data analysis

Mass spectrometry and data analysis was performed as described (Niu et al., 2019). Briefly, LC separation was performed using an Acclaim PepMap RSLC C18 capillary column (75 μm × 25 cm; 2 μm particles, 100 Å) (Thermo fisher Scientific, San Jose, CA). Gradient elution was performed using an Ultimate 3000 nanoLC system (Thermo fisher Scientific, San Jose, CA). The flow rate was set at 300 nl/min. Mobile phase A was 0.1% formic acid in water and mobile phase B was 0.1% formic acid in acetonitrile. The analytical gradient lasted for 90 min including the following steps: 1) 10 min of equilibration with 3% B; 2) the composition of solvent B was increased from 3% to 7% in 2 min, from 7% to 20% in 50 min, and from 20% to 30% in 2 min; 3) a washing and equilibration step when solvent B was increased to 80% in 1 min and was held for 8 min; 4) the composition of solvent B was returned to 3% in 0.1 min and was held for 17 min.

Analysis was performed using an Orbitrap Fusion Lumos mass spectrometer (Thermo Fisher Scientific, San Jose, CA) operating in positive ion mode. The ESI spray voltage was set at 2300 KV and the ion transfer tube temperature was set at 300°C. MS and MS/MS scans were performed using high resolution Orbitrap, with resolution at 60K and 15K, respectively. Data-dependent acquisition (DDA) mode was performed with a cycle time of 3 s. The mass range of the full MS scan defines m/z 400 to 1600, and in MS/MS starts from m/z 110. The collision energy was set at 30%. Three biological repeats of each sample were performed. Proteome Discoverer 2.2.0.388 was used for protein identification and quantification with the following settings: (1) fixed modification: cysteine carbamidomethylation (+57.021 Da); (2) dynamic modification: methionine oxidation (+15.995 Da) and acetylation (+42.011) at the N terminus of the protein; (3) trypsin was used for digestion with one missed cleavage allowed; (4) peptide ion tolerance: 10 ppm; (5) fragment ion tolerance: 20 ppm; (6) the protein sequence database of *Homo sapiens* was downloaded from Uniprot (updated 11-2018) for database searching and identification with a false discovery rate (FDR) <0.01 considered reliable; (7) the minora algorithm based label-free quantification (LFQ) was performed using the ratio of the intensity of precursor ions; (8) unique and razor peptides of proteins were selected for protein relative quantification; (9) Normalization to the protein median of each sample was used to correct experimental bias based on the total peptide amount.

The following criteria was used for selecting the identified proteins for quantitative analysis: 1) proteins identified with two or more unique peptides; 2) proteins which were successfully quantified in all of the three biological repeats of at least one of the experimental groups were selected for further analysis. The abundance ratio of each identified proteins was determined using pairwise ratio based calculation where the median value of the abundance ratio of all matched peptides from three biological repeats was used as the abundance ratio of the identified protein. The maximum allowed fold change was set as 100. Student’s t-test was used to compare the significant changes between two experimental groups based on the abundance values of the identified protein in three biological repeats in the two experimental groups.

## Acknowledgements

We thank Dr. Randy Schekman (UC Berkeley) to provide purified COPII components. We thank Dr. Xiao-wei Chen (Peking University, China) to provide antibodies against SURF4. This work was supported by the Hong Kong Research Grants Council Grants 26100315, 16101116, 16102218, 16103319, AoE/M-05/12, C4002-17G and C4012-16E to Y.G. This work was also supported by a grant from National Natural Science Foundation of China (NSFC 31871421 to Y.G. and NSFC 81601828 to H.Y.) and a free explore project from Shenzhen Science and Technology Innovation Committee (JCYJ20180306174847511) to Y.G. In addition, this work was supported by Medical Research Council (MC_UP_1201/10) to EAM. Z.Y. is supported by RGC CRF Grant No. C5031-14E and the University Research Facility in Chemical and Environmental Analysis of PolyU. LJ is supported by grants from RGC of HK (C4012-16E, C4002-17G and AoE/M-05/12).

---

**Figure S1.**
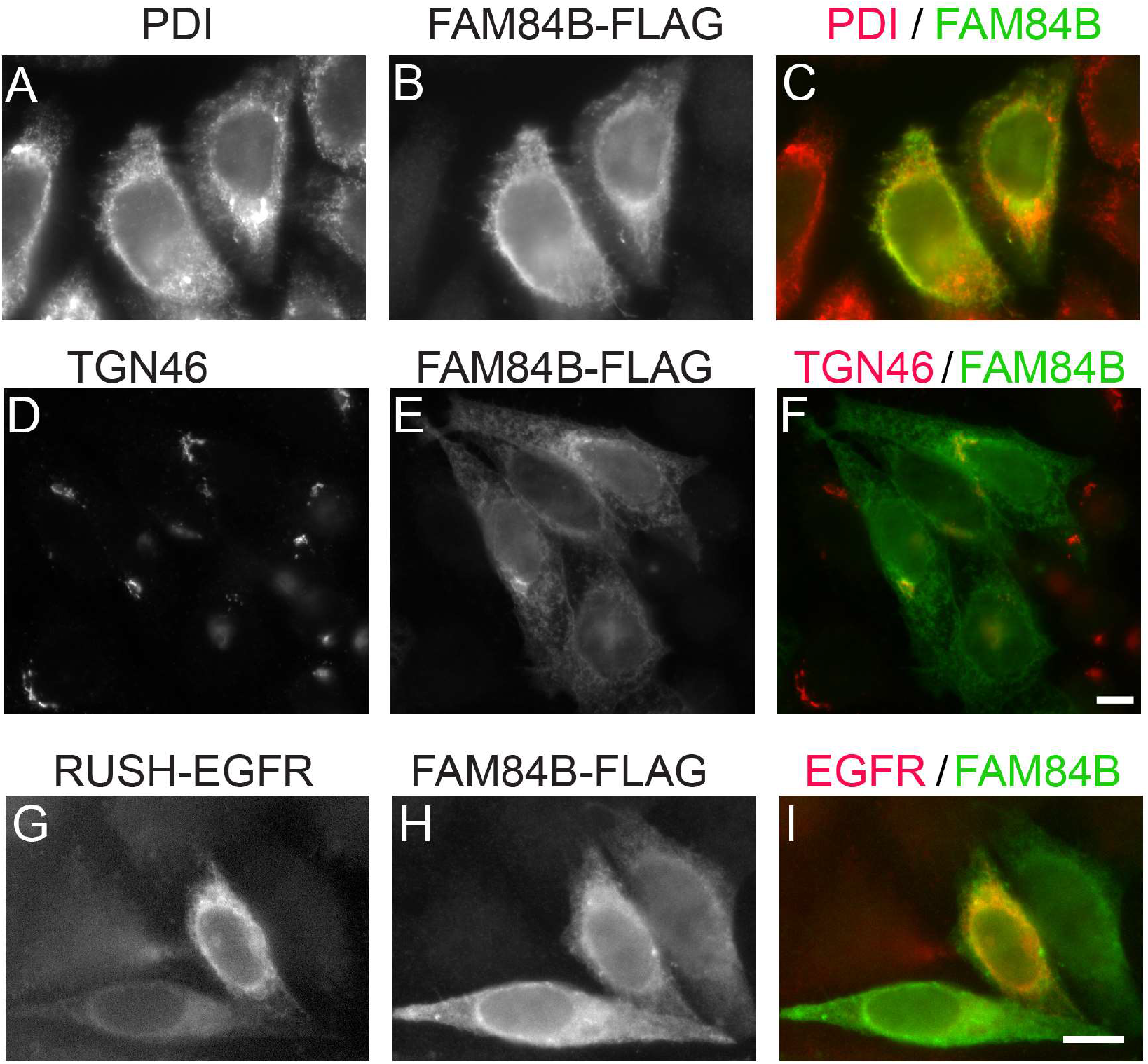
Localization analysis of FAM84B-FLAG. HeLa cells were transfected with FAM84B-FLAG (A-F) or co-transfected with FAM84B-FLAG and RUSH-EGFR (G-I). Day 1 after transfection, the localizations of the indicated proteins were analyzed by immunofluorescence. *Scale bar*, 10 μm.

**Figure S2.**
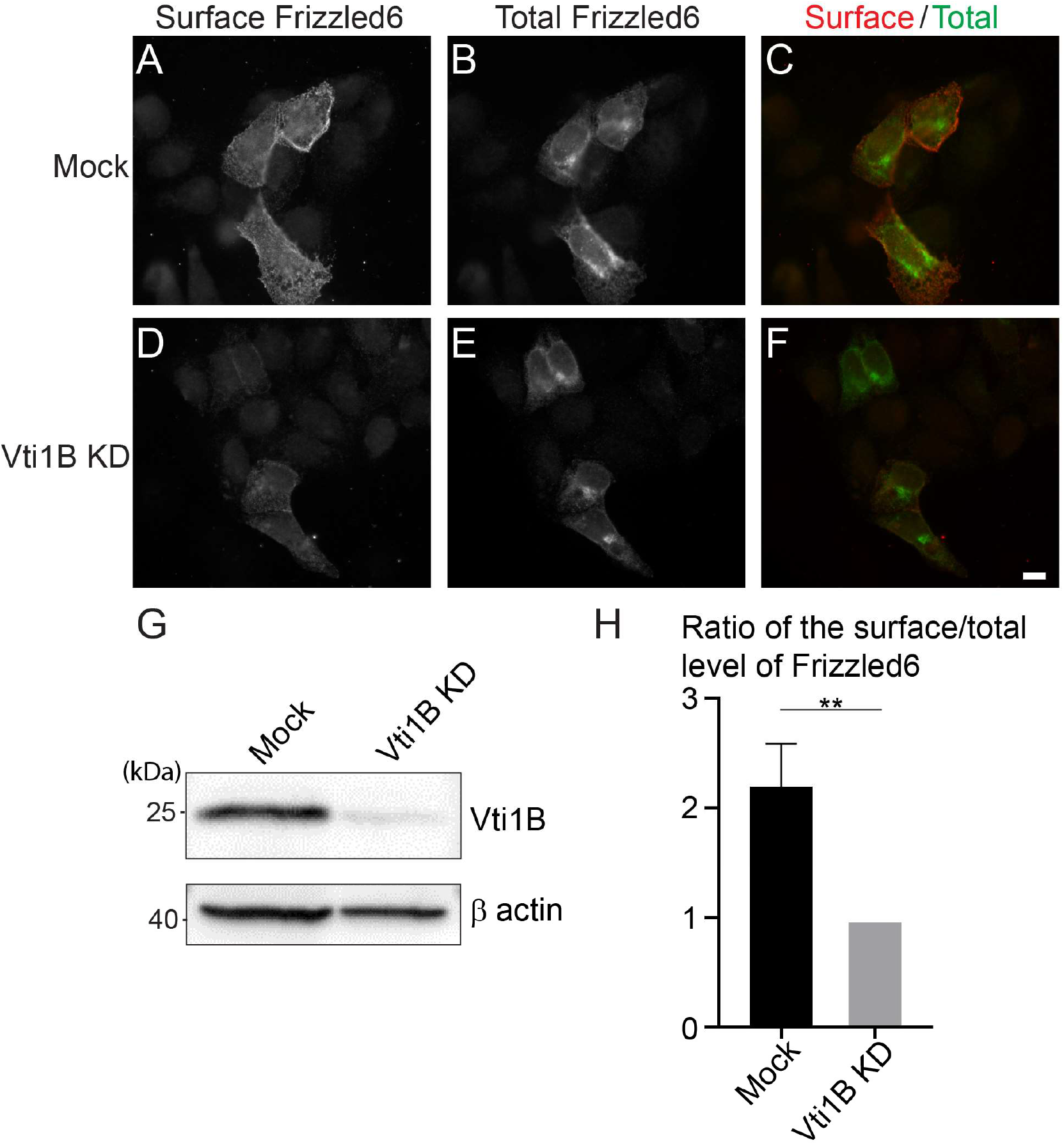
Vti1B regulates TGN-to-cell surface delivery of Frizzled6. **A-F.** HeLa cells were mock transfected (A–C) or transfected with siRNA against Vti1B (D-F) and re-transfected after 48 h with plasmids encoding HA-Frizzled6. On day 3 after knockdown, cells were incubated at 20 °C for 2 h then shifted to 32 °C for 90 min. After incubation, cells were analyzed by immunofluorescence. The surface-localized HA-Frizzled6 and the total HA-Frizzled6 were stained by mouse and rabbit anti-HA antibodies, respectively. Scale bar, 10 μm. **G.** HeLa cells were mock transfected or transfected with siRNA against Vti1B. On day 3 after transfection, cells were analyzed by immunoblot. **H.** Quantification of the ratio of the fluorescent level of surface Frizzled6 over the fluorescent level of total Frizzled6 after incubation at 32 °C (mean ± S.D.; n = 3; >120 cells from 9 random imaging fields counted for each experiment). In each experiment, the total fluorescent level was normalized to that in Vti1B KD cells. **, p < 0.01.

**Table S1.** Identification of cytosolic proteins that are associated with vesicles in a GTP dependent manner and cargo proteins that are packaged into vesicles in a GTP-hydrolysis dependent manner.

**Table S2.** Identification of cargo proteins and cytosolic proteins that are dependent on Sar1A to be associated with transport vesicles

**Table S3.** Identification of cargo proteins that depend on ERGIC53 or SURF4 for packaging into transport vesicles

## References

Adolf, F., M. Rhiel, B. Hessling, Q. Gao, A. Hellwig, J. Bethune, and F.T. Wieland. 2019. Proteomic Profiling of Mammalian COPII and COPI Vesicles. Cell Rep. 26:250–265 e255.

Appenzeller, C., H. Andersson, F. Kappeler, and H.P. Hauri. 1999. The lectin ERGIC-53 is a cargo transport receptor for glycoproteins. Nat Cell Biol. 1:330–334.

Aridor, M., S.I. Bannykh, T. Rowe, and W.E. Balch. 1995. Sequential coupling between COPII and COPI vesicle coats in endoplasmic reticulum to Golgi transport. J Cell Biol. 131:875–893.

Barlowe, C., and A. Helenius. 2016. Cargo Capture and Bulk Flow in the Early Secretory Pathway. Annu Rev Cell Dev Biol. 32:197–222.

Boncompain, G., S. Divoux, N. Gareil, H. de Forges, A. Lescure, L. Latreche, V. Mercanti, F. Jollivet, G. Raposo, and F. Perez. 2012. Synchronization of secretory protein traffic in populations of cells. Nat Methods. 9:493–498.

Cheng, C., H. Cui, L. Zhang, Z. Jia, B. Song, F. Wang, Y. Li, J. Liu, P. Kong, R. Shi, Y. Bi, B. Yang, J. Wang, Z. Zhao, Y. Zhang, X. Hu, J. Yang, C. He, Z. Zhao, J. Wang, Y. Xi, E. Xu, G. Li, S. Guo, Y. Chen, X. Yang, X. Chen, J. Liang, J. Guo, X. Cheng, C. Wang, Q. Zhan, and Y. Cui. 2016. Genomic analyses reveal FAM84B and the NOTCH pathway are associated with the progression of esophageal squamous cell carcinoma. Gigascience. 5:1.

Chidambaram, S., N. Mullers, K. Wiederhold, V. Haucke, and G.F. von Mollard. 2004. Specific interaction between SNAREs and epsin N-terminal homology (ENTH) domains of epsin-related proteins in trans-Golgi network to endosome transport. J Biol Chem. 279:4175–4179.

Chidambaram, S., J. Zimmermann, and G.F. von Mollard. 2008. ENTH domain proteins are cargo adaptors for multiple SNARE proteins at the TGN endosome. J Cell Sci. 121:329–338.

Christoforidis, S., and M. Zerial. 2000. Purification and identification of novel Rab effectors using affinity chromatography. Methods. 20:403–410.

Dancourt, J., and C. Barlowe. 2010. Protein sorting receptors in the early secretory pathway. Annu Rev Biochem. 79:777–802.

Dascher, C., and W.E. Balch. 1996. Mammalian Sly1 regulates syntaxin 5 function in endoplasmic reticulum to Golgi transport. J Biol Chem. 271:15866–15869.

Emmer, B.T., G.G. Hesketh, E. Kotnik, V.T. Tang, P.J. Lascuna, J. Xiang, A.C. Gingras, X.W. Chen, and D. Ginsburg. 2018. The cargo receptor SURF4 promotes the efficient cellular secretion of PCSK9. Elife. 7.

Guo, Y., and A.D. Linstedt. 2006. COPII-Golgi protein interactions regulate COPII coat assembly and Golgi size. J Cell Biol. 174:53–63.

Guo, Y., D.W. Sirkis, and R. Schekman. 2014. Protein sorting at the trans-Golgi network. Annu Rev Cell Dev Biol. 30:169–206.

Guo, Y., G. Zanetti, and R. Schekman. 2013. A novel GTP-binding protein-adaptor protein complex responsible for export of Vangl2 from the trans Golgi network. Elife. 2:e00160.

Herzig, Y., H.J. Sharpe, Y. Elbaz, S. Munro, and M. Schuldiner. 2012. A systematic approach to pair secretory cargo receptors with their cargo suggests a mechanism for cargo selection by Erv14. PLoS Biol. 10:e1001329.

Hirst, J., S.E. Miller, M.J. Taylor, G.F. von Mollard, and M.S. Robinson. 2004. EpsinR is an adaptor for the SNARE protein Vti1b. Mol Biol Cell. 15:5593–5602.

Jiang, Y., X. Lin, A. Kapoor, L. He, F. Wei, Y. Gu, W. Mei, K. Zhao, H. Yang, and D. Tang. 2019. FAM84B promotes prostate tumorigenesis through a network alteration. Ther Adv Med Oncol. 11:1758835919846372.

Jin, H., S.R. White, T. Shida, S. Schulz, M. Aguiar, S.P. Gygi, J.F. Bazan, and M.V. Nachury. 2010. The conserved Bardet-Biedl syndrome proteins assemble a coat that traffics membrane proteins to cilia. Cell. 141:1208–1219.

Kim, J., B. Kleizen, R. Choy, G. Thinakaran, S.S. Sisodia, and R.W. Schekman. 2007. Biogenesis of gamma-secretase early in the secretory pathway. J Cell Biol. 179:951–963.

Lee, M.C., E.A. Miller, J. Goldberg, L. Orci, and R. Schekman. 2004. Bi-directional protein transport between the ER and Golgi. Annu Rev Cell Dev Biol. 20:87–123.

Ma, T., B. Li, R. Wang, P.K. Lau, Y. Huang, L. Jiang, R. Schekman, and Y. Guo. 2018. A mechanism for differential sorting of the planar cell polarity proteins Frizzled6 and Vangl2 at the trans-Golgi network. J Biol Chem. 293:8410–8427.

Mao, Y., R. Tu, Y. Huang, D. Mao, Z. Yang, P.K. Lau, J. Wang, J. Ni, Y. Guo, and T. Xie. 2019. The exocyst functions in niche cells to promote germline stem cell differentiation by directly controlling EGFR membrane trafficking. Development. 146.

Merte, J., D. Jensen, K. Wright, S. Sarsfield, Y. Wang, R. Schekman, and D.D. Ginty. 2010. Sec24b selectively sorts Vangl2 to regulate planar cell polarity during neural tube closure. Nat Cell Biol. 12:41–46; sup pp 41–48.

Miller, S.E., B.M. Collins, A.J. McCoy, M.S. Robinson, and D.J. Owen. 2007. A SNARE-adaptor interaction is a new mode of cargo recognition in clathrin-coated vesicles. Nature. 450:570–574.

Nichols, W.C., U. Seligsohn, A. Zivelin, V.H. Terry, C.E. Hertel, M.A. Wheatley, M.J. Moussalli, H.P. Hauri, N. Ciavarella, R.J. Kaufman, and D. Ginsburg. 1998. Mutations in the ER-Golgi intermediate compartment protein ERGIC-53 cause combined deficiency of coagulation factors V and VIII. Cell. 93:61–70.

Niu, L., T. Ma, F. Yang, B. Yan, X. Tang, H. Yin, Q. Wu, Y. Huang, Z.P. Yao, J. Wang, Y. Guo, and J. Hu. 2019. Atlastin-mediated membrane tethering is critical for cargo mobility and exit from the endoplasmic reticulum. Proc Natl Acad Sci U S A. 116:14029–14038.

Nogueira, C., P. Erlmann, J. Villeneuve, A.J. Santos, E. Martinez-Alonso, J.A. Martinez-Menarguez, and V. Malhotra. 2014. SLY1 and Syntaxin 18 specify a distinct pathway for procollagen VII export from the endoplasmic reticulum. Elife. 3:e02784.

Nyfeler, B., V. Reiterer, M.W. Wendeler, E. Stefan, B. Zhang, S.W. Michnick, and H.P. Hauri. 2008. Identification of ERGIC-53 as an intracellular transport receptor of alpha1-antitrypsin. J Cell Biol. 180:705–712.

Nyfeler, B., B. Zhang, D. Ginsburg, R.J. Kaufman, and H.P. Hauri. 2006. Cargo selectivity of the ERGIC-53/MCFD2 transport receptor complex. Traffic. 7:1473–1481.

Ong, Y.S., B.L. Tang, L.S. Loo, and W. Hong. 2010. p125A exists as part of the mammalian Sec13/Sec31 COPII subcomplex to facilitate ER-Golgi transport. J Cell Biol. 190:331–345.

Saegusa, K., M. Sato, N. Morooka, T. Hara, and K. Sato. 2018. SFT-4/Surf4 control ER export of soluble cargo proteins and participate in ER exit site organization. J Cell Biol. 217:2073–2085.

Shaywitz, D.A., P.J. Espenshade, R.E. Gimeno, and C.A. Kaiser. 1997. COPII subunit interactions in the assembly of the vesicle coat. J Biol Chem. 272:25413–25416.

Shugrue, C.A., E.R. Kolen, H. Peters, A. Czernik, C. Kaiser, L. Matovcik, A.L. Hubbard, and F. Gorelick. 1999. Identification of the putative mammalian orthologue of Sec31P, a component of the COPII coat. J Cell Sci. 112 (Pt 24):4547–4556.

Tse, Y.C., B. Mo, S. Hillmer, M. Zhao, S.W. Lo, D.G. Robinson, and L. Jiang. 2004. Identification of multivesicular bodies as prevacuolar compartments in Nicotiana tabacum BY-2 cells. Plant Cell. 16:672–693.

Vollenweider, F., F. Kappeler, C. Itin, and H.P. Hauri. 1998. Mistargeting of the lectin ERGIC-53 to the endoplasmic reticulum of HeLa cells impairs the secretion of a lysosomal enzyme. J Cell Biol. 142:377–389.

Wakana, Y., J. van Galen, F. Meissner, M. Scarpa, R.S. Polishchuk, M. Mann, and V. Malhotra. 2012. A new class of carriers that transport selective cargo from the trans Golgi network to the cell surface. EMBO J. 31:3976–3990.

Wang, J., M. Gossing, P. Fang, J. Zimmermann, X. Li, G.F. von Mollard, L. Niu, and M. Teng. 2011. Epsin N-terminal homology domains bind on opposite sides of two SNAREs. Proc Natl Acad Sci U S A. 108:12277–12282.

Wong, N., Y. Gu, A. Kapoor, X. Lin, D. Ojo, F. Wei, J. Yan, J. de Melo, P. Major, G. Wood, T. Aziz, J.C. Cutz, M. Bonert, A.J. Patterson, and D. Tang. 2017. Upregulation of FAM84B during prostate cancer progression. Oncotarget. 8:19218–19235.

Yin, Y., M.R. Garcia, A.J. Novak, A.M. Saunders, R.S. Ank, A.S. Nam, and L.W. Fisher. 2018. Surf4 (Erv29p) binds amino-terminal tripeptide motifs of soluble cargo proteins with different affinities, enabling prioritization of their exit from the endoplasmic reticulum. PLoS Biol. 16:e2005140.

Zhang, B., M.A. Cunningham, W.C. Nichols, J.A. Bernat, U. Seligsohn, S.W. Pipe, J.H. McVey, U. Schulte-Overberg, N.B. de Bosch, A. Ruiz-Saez, G.C. White, E.G. Tuddenham, R.J. Kaufman, and D. Ginsburg. 2003. Bleeding due to disruption of a cargo-specific ER-to-Golgi transport complex. Nat Genet. 34:220–225.

Zhang, B., R.J. Kaufman, and D. Ginsburg. 2005. LMAN1 and MCFD2 form a cargo receptor complex and interact with coagulation factor VIII in the early secretory pathway. J Biol Chem. 280:25881–25886.

